# Diversification of DNA-binding specificity via permissive and specificity-switching mutations in the ParB/Noc protein family

**DOI:** 10.1101/724823

**Authors:** Adam S. B. Jalal, Ngat T. Tran, Clare E. Stevenson, Elliot W. Chan, Rebecca Lo, Xiao Tan, Agnes Noy, David M. Lawson, Tung B. K. Le

## Abstract

Specific interactions between proteins and DNA are essential to many biological processes. Yet it remains unclear how the diversification in DNA-binding specificity was brought about, and what were the mutational paths that led to changes in specificity. Using a pair of evolutionarily related DNA-binding proteins, each with a different DNA preference (ParB and Noc: both having roles in bacterial chromosome maintenance), we show that specificity is encoded by a set of four residues at the protein-DNA interface. Combining X-ray crystallography and deep mutational scanning of the interface, we suggest that permissive mutations must be introduced before specificity-switching mutations to reprogram specificity, and that mutational paths to a new specificity do not necessarily involve dual-specificity intermediates. Overall, our results provide insight into the possible evolutionary history of ParB and Noc, and in a broader context, might be useful in understanding the evolution of other classes of DNA-binding proteins.

## INTRODUCTION

In living organisms, hundreds of DNA-binding proteins carry out a plethora of roles in homeostasis, transcriptional regulation in response to stress, and in maintenance and transmission of genetic information. These DNA-binding proteins do so faithfully due to their distinct DNA-binding specificity towards their cognate DNA sites. Yet it remains unclear how related proteins, sometimes with a very similar DNA-recognition motif, can recognize entirely different DNA sites. What were the changes at the molecular level that brought about the diversification in DNA-binding specificity? As these proteins evolved, did the intermediates in this process drastically switch DNA-binding specificity or did they transit gradually through promiscuous states that recognized multiple DNA sequences? Among many ways to evolve new biological innovations, gene duplication and neo-functionalization have been widely implicated as a major force in evolution [1–5]. In this process, after a gene was duplicated, one copy retained the original function while the other accumulated beneficial and diverging mutations that produced a different protein with a new function. In the case of DNA-binding proteins, a new function could be the recognition of an entirely different DNA site. In this work, we employed a pair of related DNA-binding proteins (ParB and Noc) that are crucial for bacterial chromosome segregation and maintenance to better understand factors that might have influenced the evolution of a new DNA-binding specificity.

ParB (<u>Par</u>titioning Protein <u>B</u>) is important for faithful chromosome segregation in two-thirds of bacterial species [6,7]. The centromere-like *parS* DNA locus is the first to be segregated following chromosome replication [6–10]. *parS* is bound by ParB, which in turn interacts with ParA and SMC proteins to partition the ParB-*parS* nucleoprotein complex, hence the chromosome, into each daughter cell [7,11–18] (Figure 1A). ParB specifically recognizes and binds to *parS*, a palindromic sequence (Figure 1A) that can be present as multiple copies on the bacterial chromosome but almost always locate close to the origin of replication (*oriC*) on each chromosome (Figure 1A) [6–8,17,19–22]. ParB proteins are widely distributed in bacteria and so must have appeared early in evolution (Figure 1B) [6].

**Figure 1.**
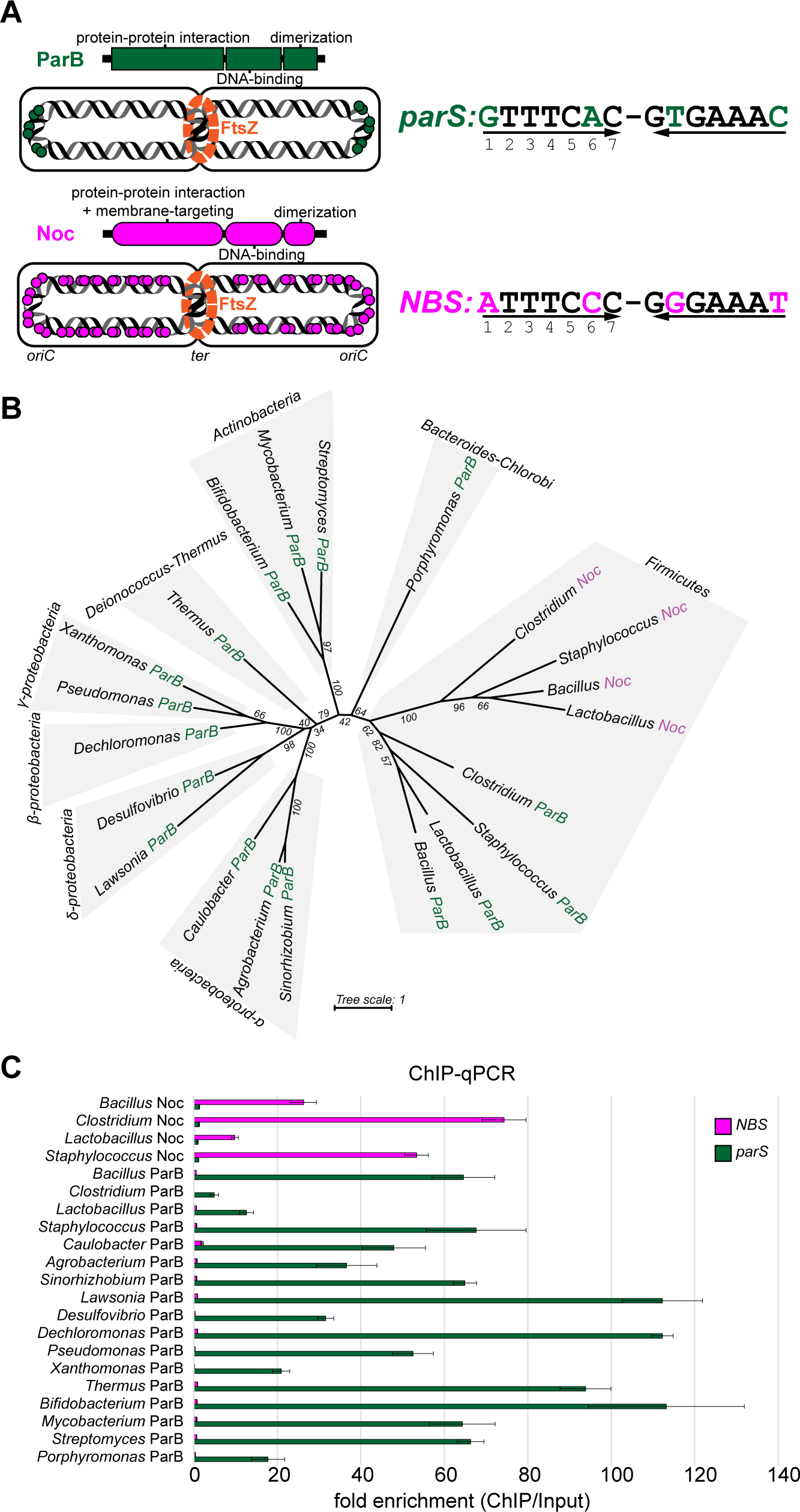
DNA-binding specificity for *parS* and *NBS* is conserved among ParB and Noc orthologs. **(A)** The domain architecture of ParB (dark green) and Noc (magenta) together with their respective cognate DNA-binding sites, *parS* and *NBS*. Sequence differences between *parS* and *NBS* are highlighted (*parS*: dark green, *NBS*: magenta). The genome-wide distributions of *parS* and *NBS* sites (dark green and magenta circles, respectively) are also shown schematically. **(B)** An unrooted maximum likelihood tree that shows the restrictive distribution of Noc orthologs (magenta branches) to the Firmicutes clade. Bootstrap support values are shown for branches. **(C)** The *in vivo* binding preferences of ParB/Noc to *parS*/*NBS* as measured by ChIP-qPCR. Error bars represent standard deviation (SD) from three replicates. An *E. coli* strain with a single *parS* and *NBS* site engineered onto the chromosome was used as a heterologous host for the expression of FLAG-tagged ParB/Noc.

Noc (<u>N</u>ucleoid <u>Oc</u>clusion Factor), a ParB-related protein, was first discovered in *Bacillus subtilis* [23,24]. Like ParB, Noc has a three-domain architecture: an N-terminal domain for protein-protein interaction and for targeting Noc to the cell membrane, a central DNA-binding domain, and a C-terminal dimerization domain [24,25] (Figure 1A). In contrast to ParB, Noc recognizes a DNA-binding sequence called *NBS* (<u>N</u>oc <u>B</u>inding <u>S</u>ite) [25,26] (Figure 1A). The role of Noc is also different from ParB; Noc functions to prevent the cell division machinery from assembling in the vicinity of the nucleoid, which might be otherwise guillotined, thereby damaging the DNA[24,25] (Figure 1B). In other words, Noc has a role in preserving the integrity of the chromosome. The genome-wide distribution of *NBS* is also drastically different from that of *parS*. While *parS* sites are restricted in the region around *oriC, NBS* distributes widely on the genome, except near the terminus of replication (*ter*) [25,26]. The absence of *NBS* near *ter* is crucial to direct the formation of the FtsZ ring and cell division to mid-cell (Figure 1A). Because of their genomic proximity (Figure S1) and high sequence similarity, it was suggested that *noc* resulted from a gene duplication event from *parB* [23,27]. A phylogenetic tree showed that *parB* genes are widely distributed in bacteria but *noc* genes are confined to the Firmicutes clade [27] (Figure 1B). This phylogenetic distribution is most consistent with *parB* appearing early in evolution, possibly before the split between Gram-positive and Gram-negative bacteria, and that the occurrence of *noc* is a later event that happened only in Firmicutes [27].

Here, we systematically measure the binding preferences of 17 ParB and four Noc family members to *parS* and *NBS* and find that their interactions are specific and conserved among bacterial species. We show that specificity to *parS* or *NBS* is encoded by a small set of four residues at the protein-DNA interface, and mutations in these residues are enough to reprogram DNA-binding specificity. Combining X-ray crystallography and systematic scanning mutagenesis, we show that both permissive and specificity-switching substitutions are required to acquire a new DNA-binding specificity. Guided by these findings, we generate a saturated library with ∼10^5^ variants of the specificity-defining residues in ParB, and select for mutants that bind to *parS, NBS*, or both. We discover multiple alternative combinations of residues that are capable of binding to *parS* or *NBS*. By analyzing the connectivity of functional variants in the sequence space, we suggest that permissive and specificity-switching mutations, at least when considering the four mutations in this work, must be introduced in an orderly manner to evolve a new protein-DNA interface.

## RESULTS

### DNA-binding specificity for *parS* and *NBS* is conserved within ParB and Noc family

To test whether ParB and Noc family members retained their DNA-binding specificity, we selected a group of 17 ParB and four Noc from various bacterial clades for characterization (Figure 1B and Figure S1A). ParB or Noc proteins were expressed individually in *Escherichia coli* and were engineered with an N-terminal FLAG tag for immunoprecipitation. We performed ChIP-qPCR and ChIP-seq experiments to quantify the level of ParB or Noc that are bound at a single *parS* or *NBS* site engineered onto the *E. coli* chromosome (Figure 1C and Figure S1B). *E. coli* is a perfect heterologous host for this experiment as it does not possess native ParB/Noc homologs and there are no *parS*/*NBS* sites in its genome. As shown in Figure 1C, all tested ParB proteins bind preferentially to *parS* over *NBS*, while Noc proteins prefer *NBS* to *parS*. This conservation of DNA preference suggests that there exists a set of conserved residues within each protein family (ParB or Noc) that dictate specificity.

### The co-crystal structure of the DNA-binding domain of ParB with *parS* reveals residues that contact DNA

As the first step in identifying specificity residues, we solved a 2.4 Å resolution co-crystal structure of the DNA-binding domain (DBD) of *Caulobacter crescentus* ParB bound to a 20-bp *parS* DNA duplex (Figure 2A). In the crystallographic asymmetric unit, two very similar ParB DBD monomers (RMSD = 0.1 Å) bind in a two-fold symmetric fashion to a full-size *parS* DNA duplex (Figure 2A). This structure reveals several regions of each DBD that contact *parS* (Figure 2B). Firstly, the recognition helix α4 of the helix-turn-helix motif inserts into the major grooves of the palindromic *parS* site (Figure 2B). Second, helices α6 and α8 contribute residues to the protein-DNA interface (Figure 2B). Lastly, several lysine and arginine residues in the loop spanning res. 236-254 contact the minor groove side of *parS* in an adjacent complex in the crystal (Figure 2A). From the structure of the complex, we identified residues that make specific contacts with the DNA bases as well as non-specific contacts with the phosphate backbone (Figure 2C). We verified the protein-DNA contacts by individually mutating each residue to alanine (Figure 2D). We found that most of the crucial residues for binding to *parS* are within the 162-234 region (Figure 2D), suggesting their importance in recognizing DNA specifically. We reasoned that specificity residues for *parS* (and *NBS*) must localize within this amino acid region in ParB (and in an equivalent region in Noc).

**Figure 2.**
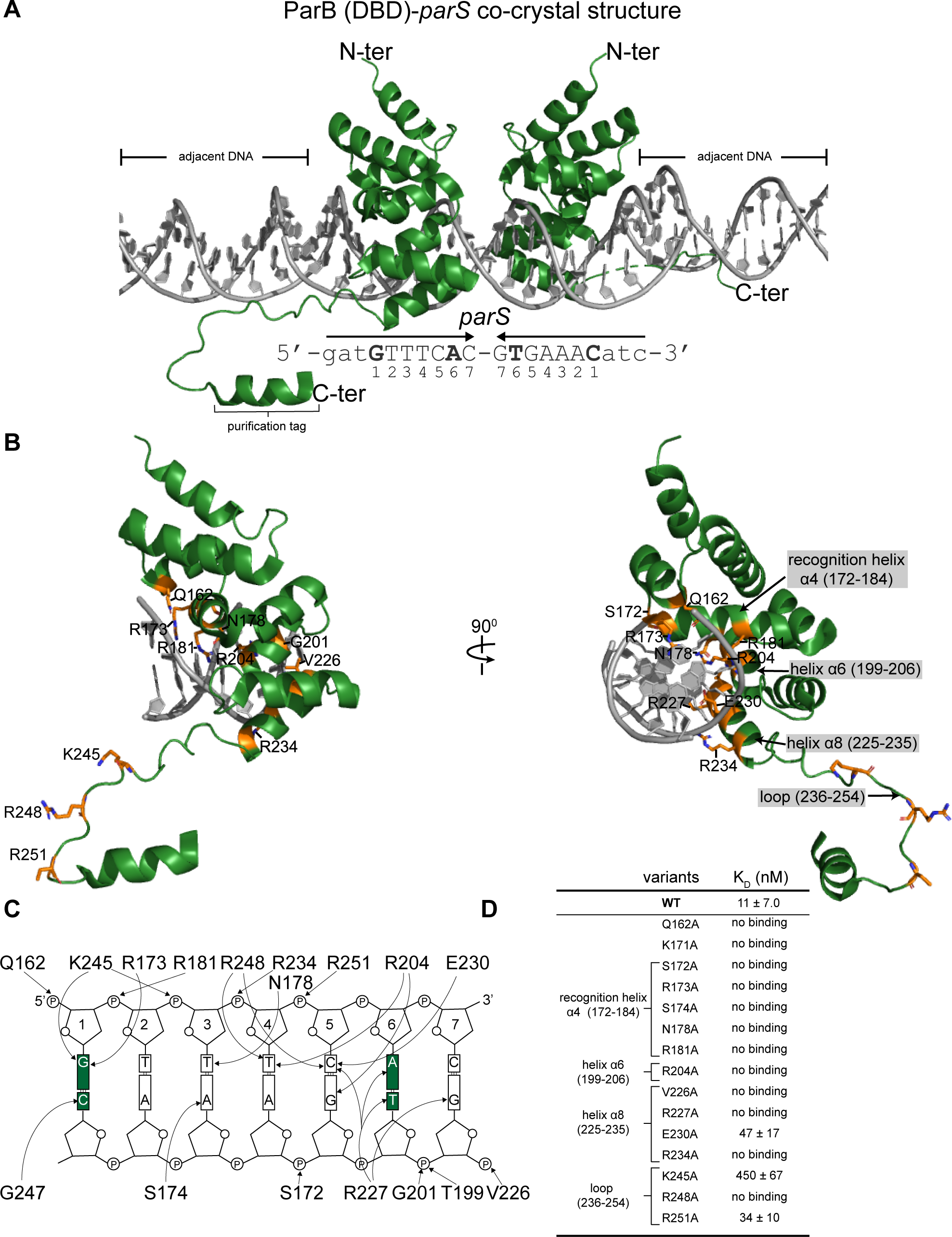
Co-crystal structure of the DNA-binding domain (DBD) of *Caulobacter* ParB with *parS*. The 2.4 Å resolution structure of two ParB (DBD) monomers (dark green) in complex with a 20-bp *parS* DNA (grey). The nucleotide sequence of the 20-bp *parS* is shown below the structure; bases (Guanine 1 and Adenine 6) that are different from *NBS* are in bold. The purification tag is also visible in one of the DBD monomers. Loop (236-254) contacts the adjacent DNA in the crystal lattice. **(B)** One monomer of ParB (DBD) is shown in complex with a *parS* half-site; residues that contact the DNA are labeled and colored in orange. **(C)** Schematic representation of ParB (DBD)-*parS* interactions. For simplicity, only a *parS* half-site is shown. The two bases at position 1 and 6 that are different between *parS* and *NBS* are highlighted in dark green. **(D)** Alanine scanning mutagenesis and the *in vitro* binding affinities (K_D_) ± SD of ParB variants to *parS* DNA.

### Mutations at four residues at the ParB-*parS* interface are sufficient to reprogram DNA-binding specificity towards *NBS*

To discover the region of Noc that determines specificity for *NBS*, we constructed a series of chimeric proteins in which different regions of *Caulobacter* ParB were replaced with the corresponding regions of *B. subtilis* Noc (Figure 3A). Replacing the entire region (res. 162-230) containing the helix-turn-helix motif, helix α6, and part of helix α8 with the corresponding region of *B. subtilis* Noc produced a chimera that binds to both *parS* and *NBS*, but with a preference for *NBS* (Chimera 1, Figure 3A). Swapping a smaller region (res. 162-207) containing just the helix-turn-helix motif and an adjacent helix α6 created a chimera that has an improved specificity for *NBS*, albeit with a lower binding affinity (Chimera 4, Figure 3A). These results suggest that the region (res. 162-207) might contain the core set of specificity residues for *NBS*.

**Figure 3.**
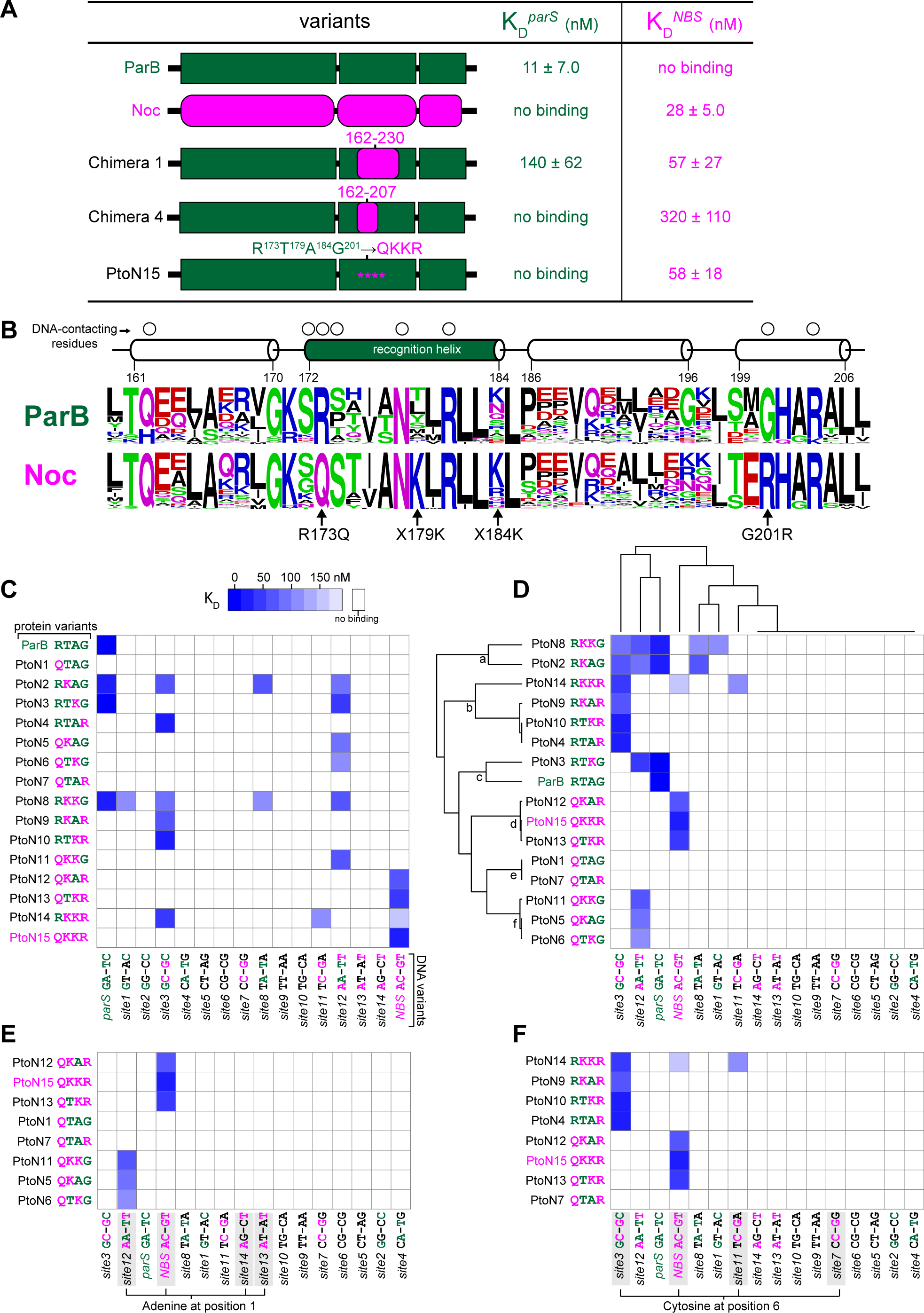
Mutations at four residues at the ParB-*parS* interface are sufficient to reprogram DNA-binding specificity towards *NBS.* **(A)** Mutations in a subset of residues in the region between res. 162-207 (ParB’s numbering) can reprogram interaction specificity. ParB (or segments of amino acids from ParB) and Noc (or equivalent segment in Noc) are shown in dark green and magenta, respectively. The affinity of protein-DNA interaction was expressed as dissociation constant (K_D_) ± SD. **(B)** The sequence alignment of ParB (∼1800 sequences) and Noc (∼400 sequences) orthologs. Amino acids are colored based on their chemical properties (GSTYC: polar; QN: neutral; KRH: basic; DE: acidic; and AVLIPWFM: hydrophobic). The secondary structure of the amino acid region (res. 162-207) is shown above the sequence alignment, together with residues (open circles) that contact DNA in the ParB (DBD)-*parS* structure (Figure 2). **(C)** Systematic scanning mutagenesis of the protein-DNA interface reveals the contribution of each specificity residue to the DNA-binding preference. Interactions between ParB + 15 PtoN intermediates with 16 DNA sites are represented as a heatmap where each matrix position reflects a K_D_ value. Amino acid residues/bases from ParB/*parS* are colored in dark green, and those from Noc/*NBS* in magenta. **(D)** A hierarchical clustering of data in panel **C** in both protein and DNA dimensions. **(E)** A simplified heatmap where only PtoN intermediates with a glutamine (Q) at position 173 are shown. **(F)** A simplified heatmap where only PtoN intermediates with an arginine (R) at position 201 are shown.

To better understand the high degree of specificity conserved within the ParB and Noc families, we mapped a sequence alignment of ∼1800 ParB and ∼400 Noc orthologs onto the ParB (DBD)-*parS* crystal structure to determine amino acid sequence preferences for those residues required for interaction specificity (Figure 3B). We focused our attention on the region between residues 162 and 207, which was shown above to contain the core specificity residues (Figure 3B). Of those amino acids that contact *parS* (Figure 2B-C), six residues (Q162, G170, K171, S172, N178, and R204) are conserved between ParB and Noc family members (Figure 3B). Two residues (R173 and G201) in ParB that contact *parS* but are changed to Q173 and R201, respectively, in Noc homologs (Figure 3B). Other residues at positions 179 and 184 vary among ParB homologs but are almost invariably a lysine in Noc family members (Figure 3B). We hypothesized that these amino acids (Q173, K179, K184, and R201) (Figure 3B) are specificity residues that dictate Noc preference for *NBS*. To test this hypothesis, we generated a variant of *Caulobacter* ParB in which these four residues were introduced at the structurally equivalent positions (R173Q, T179K, A184K and G201R). We purified and tested this variant in a bio-layer interferometry assay with *parS* and *NBS*. As shown in Figure 3A, ParB (RTAG→QKKR) (PtoN15) variant completely switched its binding preference to a non-cognate *NBS* site. Hence, a core set of four residues are enough to reprogram specificity.

### Systematic dissection of ParB-*parS* and Noc-*NBS* interfaces reveals the contribution of each specificity residue to the DNA-binding preference

To systematically dissect the role of each specificity residue, we constructed a complete set of ParB mutants that have either a single, double, or triple amino acid changes between the four specificity positions, from a *parS*-preferred *Caulobacter* ParB (R^173^T^179^A^184^G^201^) to an *NBS*-preferred variant (Q^173^K^179^K^184^R^201^). We named them <u>P</u>arB-<u>to</u>-<u>N</u>oc intermediates (PtoN, 15 variants in total). To simplify the nomenclature, we named the mutants based on the specificity residues being considered, for example, an *NBS*-preferred variant (Q^173^K^179^K^184^R^201^) is shortened to PtoN15 (QKKR). ParB and 15 PtoN variants were purified and tested with a series of 16 different DNA sites, each representing a transitional state from *parS* to *NBS* with each of the two variable positions (1 and 6) changed to any of other four DNA bases (Figure 3C). We visualized 16×16 interactions as a heatmap where each matrix position reflects a dissociation constant (K_D_).

This systematic pairwise interaction screen led to several notable observations (Figure 3C). First, there are two non-functional variants (PtoN1: QTAG and PtoN7: QTAR) that were unable to interact with any of the 16 DNA sites (Figure 3C). Second, six variants (PtoN4: RTAR, PtoN5: QKAG, PtoN6: QTKG, PtoN9: RKAR, PtoN10: RTKR, and PtoN11: QKKG) switched their specificity to a DNA site that has features borrowed from both *parS* and *NBS*. Meanwhile, four variants (PtoN2: RKAG, PtoN3: RTKG, PtoN8: RKKG, PtoN14: RKKR) were promiscuous i.e. binding to multiple different DNA sites (Figure 3C). We noted that functional PtoN variants have a lysine at either position 179, 184, or both. This observation became even clearer after we performed hierarchical clustering of the interaction profile in both the protein and the DNA dimensions (Figure 3D). A single lysine at either position 179 or 184 is enough to license the DNA-binding capability to PtoN variants (nodes a, b, d, and f on the clustering tree, Figure 3D), while PtoN1 (QTAG) and PtoN7 (QTAR) that do not possess any lysine at 179/184 are non-functional (node e, Figure 3D). We suggest that K179/184 has a permissive effect that might permit Q173 and R201 to contact DNA.

Next, we wondered which base of the *NBS* site that Q173 might contact specifically. To find out, we clustered only PtoN variants that share the Q amino acid at position 173 (Figure 3E). We discovered that those variants preferred DNA sites that possess an Adenine at position 1 (Figure 3E). We applied the same approach to find the base that residue R201 might contact (Figure 3F). The emerging trend is that PtoN variants that share an R amino acid at 201 preferred DNA sites with a Cytosine at position 6 (Figure 3F). Taken together, our results suggest a model in which each specificity residue has a distinct role: Q173 recognizes Adenine 1, R201 recognizes Cytosine 6, but they can only do so in the presence of a permissive K at either position 179 or 184, or both. In the next section, we used X-ray crystallography to provide evidence to support this model.

### Co-crystal structure of the DNA-binding domain of Noc with *NBS* reveals the contribution of specificity residues to the DNA-binding preference

To understand the biophysical mechanism underlying the specificity to *NBS*, we solved the co-crystal structure of *B. subtilis* Noc (DBD) with a 22-bp *NBS* DNA duplex (Figure S2). The diffraction of the Noc (DBD)-*NBS* crystal was anisotropic. Hence, despite the 2.23 Å resolution limit, because of low completeness in the higher resolution shells resulting from the anisotropic cutoff, the resultant electron density has the appearance of lower resolution maps, approx. 3 Å resolution (Table S4 and Methods). By superimposing the structures of ParB (DBD)-*parS* and Noc (DBD)-*NBS* complexes, we observed several changes in both the protein and the DNA sites that enabled specific interactions (Figure 4). First, R173 in ParB hydrogen bonds with *parS* Guanine 1, but the shorter side chain of a corresponding Q158 in Noc is unable to bond with Guanine 1 (Figure 4A). However, a corresponding base in *NBS* (Adenine 1) positions itself closer to enable hydrogen bonding with this Q173 residue (Figure 4A); this is possibly due to conformational changes in the *NBS* site that narrows the minor groove width at the Adenine 1:Thymine −1 position (from ∼7.7 to ∼3.7 Å, Figure S3). The switch from R to Q serves to eliminate the ability of ParB to contact *parS* Guanine 1 while simultaneously establishing a new contact with *NBS* Adenine 1. The second notable changes between the two co-crystal structures occurs at position 201 (Figure 4B). G201 from ParB has no side chain, hence cannot contact Thymine −6 specifically (Figure 4B). However, the equivalent residue R186 in Noc readily forms hydrogen bonds with Guanine −6 (Figure 4B). We also observed DNA unwinding that increased both the minor and the major groove widths at the Cytosine 6:Guanine −6 position of *NBS* (from ∼7.1 to ∼8.1 Å, and from ∼10.5 to ∼11.8 Å, respectively), possibly to move Guanine −6 outwards to accommodate a longer side chain of arginine (Figure S3). The NH group in the main chain of both G201 (ParB (DBD)-*parS* structure) and R186 (Noc(DBD)-*NBS* structure) also contact DNA non-specifically via their interaction with the phosphate groups of Thymine −6 (*parS*) and Guanine −6 (*NBS*), respectively (Figure 4, see also Figure 2C and Figure S2D).

**Figure 4.**
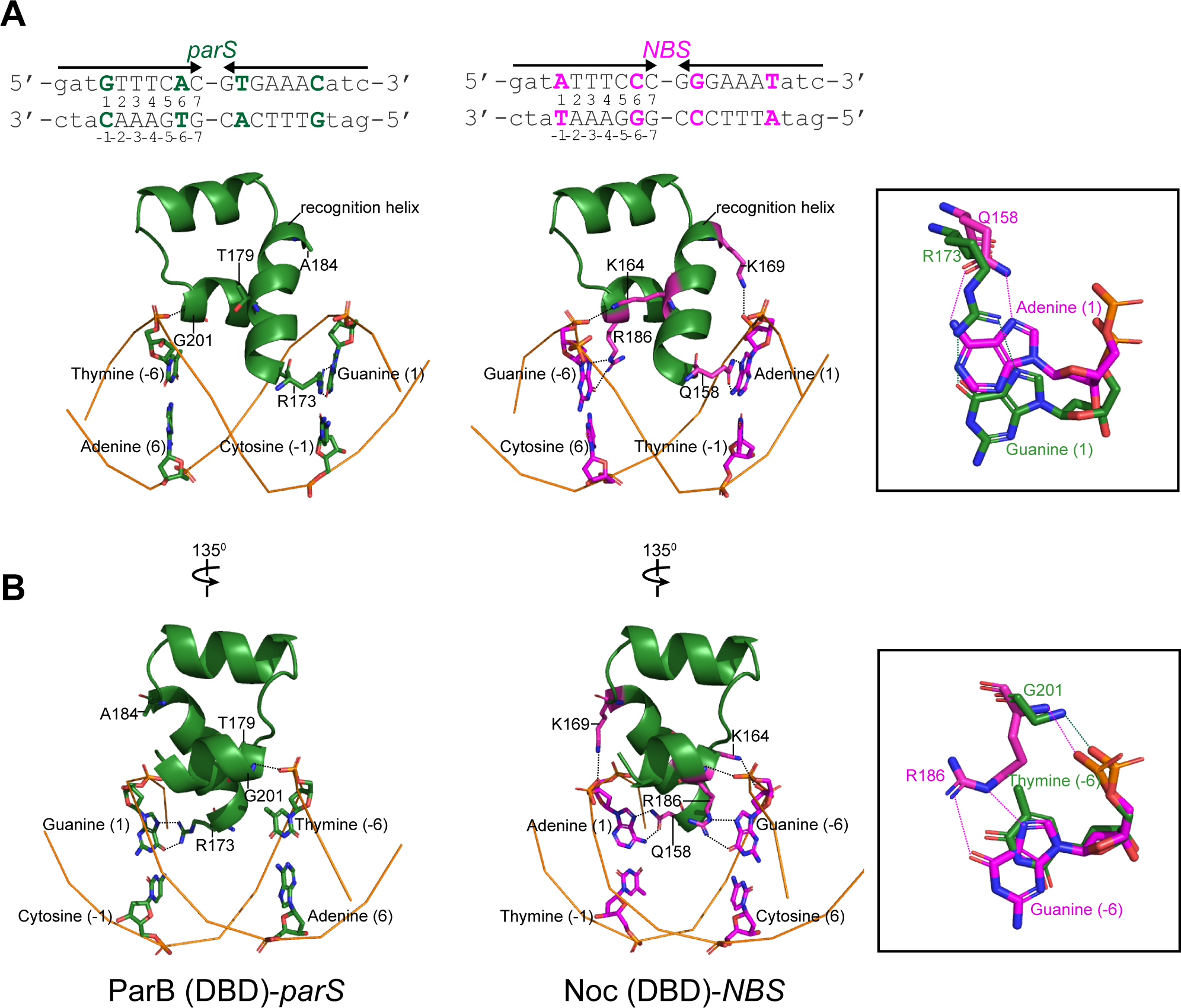
Superimposition of the ParB (DBD)-*parS* structure on the Noc (DBD)-*NBS* structure reveals the contribution of specificity residues to *NBS* binding. To simplify and highlight the roles of specificity residues, only the side chains of specificity residues and their contacting bases are shown. The amino acid regions (173-207 in ParB and the corresponding 158-192 in Noc) and the DNA backbones are shown in cartoon representation. DNA bases are numbered according to their respective positions on *parS/NBS* site. The insets show interactions between either **(A)** R173 (ParB’s numbering) and Q158 (Noc’s numbering) or **(B)** G201 (ParB’s numbering) and R186 (Noc’s numbering) and with their corresponding bases on *parS/NBS*. The side chains of K164 and K169 in Noc (DBD)-*NBS* structure contact the phosphate groups of Guanine (−5) and Thymine (2) of *NBS*, respectively (See also Figure S2D). Only the phosphate groups of Guanine (−5) and Thymine (2) in *NBS* are shown.

Our Noc (DBD)-*NBS* structure also shows the side chains of K164 and K169 make hydrogen bonds with the phosphate groups of Guanine −5 and Thymine 2 of *NBS* rather than contacting any bases specifically (Figure S2D). Lastly, molecular dynamics simulations using the Noc (DBD)-*NBS* structure as initial coordinates also suggested that side chains of K164 and K169 make hydrogen bonds or salt bridges with the DNA backbone, especially when water-mediated contacts were also considered (bonding for >99% of the whole simulation, see also Methods). The most parsimonious explanation for the permissive capability of K164/169 is that they increase DNA-binding affinity non-specifically to overcome the initial energy barrier and permit specific base contacts from Q158 and R186. Overall, our co-crystal structures are consistent with data from the systematic scanning mutagenesis.

### A high throughput bacterial one-hybrid selection reveals multiple combinations of specificity residues that enable *parS* and *NBS* recognition

While the results from our systematic scanning mutagenesis and X-ray crystallography revealed how specificity changed as individual substitutions were introduced, presumably more variety of amino acids has been sampled by Nature than those presented at the start (RTAG) and endpoint (QKKR). What are the paths, and are there many, to convert a *parS*-binding protein to an *NBS*-preferred one? Does the order of amino acid substitutions matter? To answer these questions, we explored the entire sequence space at the four specificity residues by generating a combinatorial library of ParB where positions 173, 179, 184, and 201 can be any amino acid (20^4^ or 160,000 variants lacking stop codons). We optimized a bacterial one-hybrid (B1H) assay [28] that is based on transcriptional activation of an imidazoleglycerol-phosphate dehydratase encoding gene HIS3 to enable a selection for *parS* or *NBS*-binding variants (Figure 5A and Figure S4). ParB variants were fused at their N-termini to the omega subunit of bacterial RNA polymerase. NNS codons (where N = any nucleotide and S = Cytosine or Guanine) were used to randomize the four specificity residues. All ParB variants were also engineered to contain an additional invariable mutation in the N-terminal domain (R104A) that makes ParB unable to spread [17,29] (Figure 5A). The R104A mutation does not affect the site-specific binding but enables a simpler design of the selection system by converting ParB to a conventional site-specific transcriptional activator (Figure 5A). If a ParB variant binds to a *parS* or *NBS* site engineered upstream of HIS3, it will recruit RNA polymerase to activate HIS3 expression, thereby enabling a histidine-auxotrophic *E. coli* host to survive on a minimal medium lacking histidine (Figure 5A and Figure S4). Deep sequencing of starting libraries revealed that >94% of the predicted variants were represented by at least 10 reads (Figure S5A-B), and that libraries prepared on different days were reproducible (*R*^*2*^>0.90, Figure S5C).

**Figure 5.**
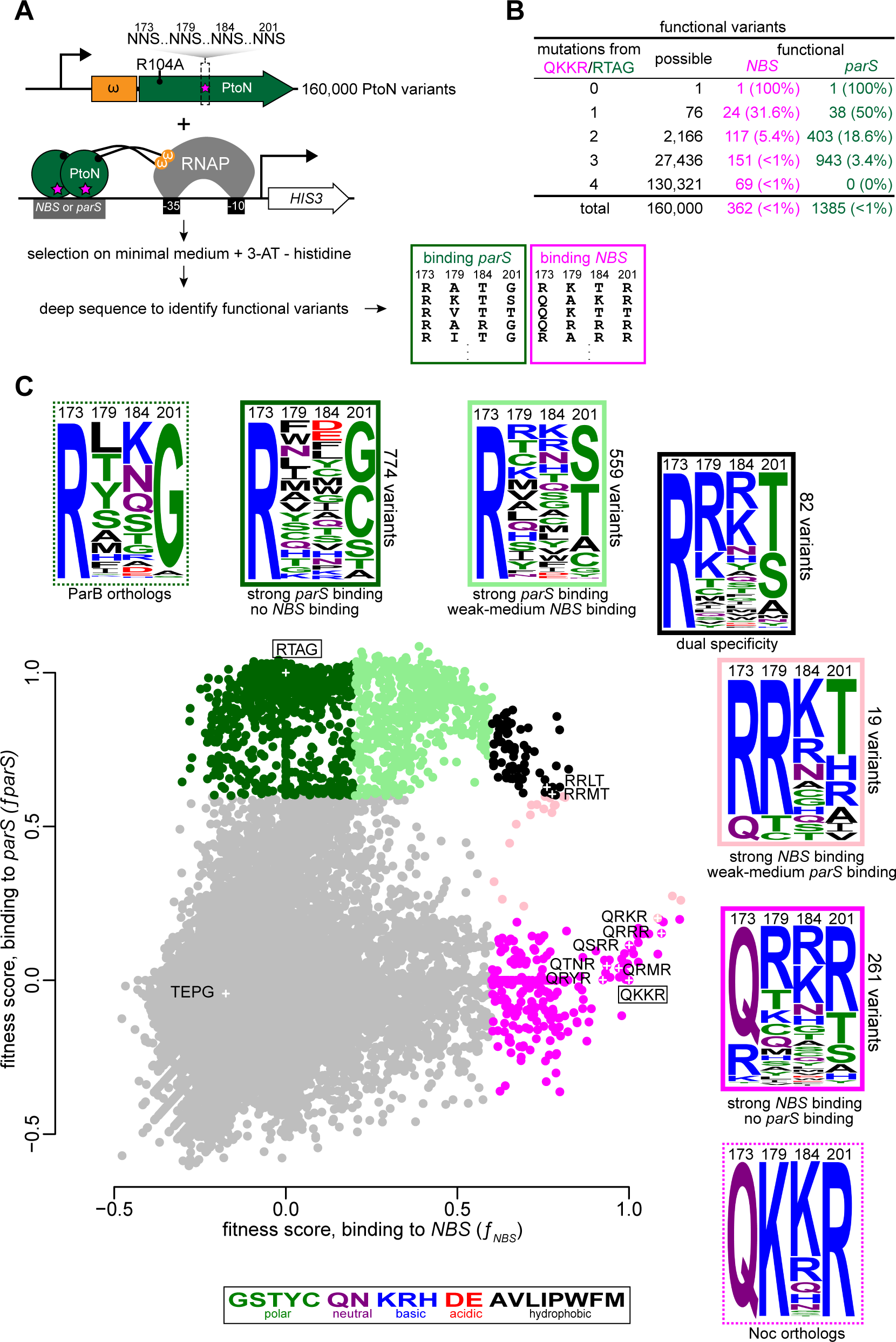
High-throughput mapping of the fitness of protein-DNA interface mutants. **(A)** The principle and design of the deep mutational scanning experiment which was based on bacterial one-hybrid assay and high-throughput sequencing. **(B)** Summary of functional *NBS*-binding and *parS*-binding variants. **(C)** Fitness scores of variants as assessed by their ability to bind *NBS* (x-axis) or *parS* (y-axis). Dark green: strong *parS* binding, no *NBS* binding (fitness score: *f*_*parS*_≥0.6, *f*_*NBS*_≤0.2); light green: strong *parS* binding, weak-to-medium *NBS* binding (*f*_*parS*_≥0.6, 0.2≤*f*_*NBS*_≤0.6); magenta: strong *NBS* binding, no *parS* binding (*f*_*NBS*_≥0.6, *f*_*parS*_≤0.2); pink: strong *NBS* binding, weak-to-medium *parS* binding (*f*_*NBS*_≥0.6, 0.2≤*f*_*parS*_≤0.6); black: dual specificity (*f*_*NBS*_≥0.6, *f*_*parS*_≥0.6). Frequency logos of each class of variants are shown together with ones for ParB/Noc orthologs. Amino acids are colored according to their chemical properties. The positions of WT ParB (RTAG), Noc (QKKR), and nine selected variants for an independent validation are also shown and labeled on the scatterplot.

To assess the ability of each ParB variant to bind to *parS* or *NBS*, we deep sequenced the relevant region on *parB* variants pre- and post-selection to reveal the underlying sequences and their abundance (Figure 5A and Figure S5C). As the strength of protein-DNA interaction is directly related to the amount of histidine being produced [28], we quantified the fitness of each variant to rank them (Figure 5C). We found 1385 and 362 variants that show strong binding to *parS* and *NBS*, respectively (Figure 5B). We then selected and verified nine variants that either bind *NBS, parS*, or both (Figure 5C) by a pairwise B1H assay and by bio-layer interferometry assay with purified proteins (Figure S6). To systematically probe the sequence space, we generated a scatter plot of ParB variant fitness when screened for binding to *parS* or *NBS* (Figure 5C). Of 362 variants that bind *NBS* strongly: 261 are *NBS* specific (i.e. no *parS* binding, magenta box), 19 show strong *NBS* binding but weak-to-medium *parS* binding (pink box), and 82 dual-specificity variants that bind both *parS* and *NBS* (black box) (Figure 5C). By comparing sequence logos, we observed that *NBS*-specific variants (magenta box) have a high proportion of the Q residue at position 173 but R is allowed; position 201 is dominantly R but polar residues (T and S) are allowed; and positively charged R and K prevail at positions 179 and 184 (Figure 5C). This sequence logo shares some features with Noc orthologs (dashed magenta box, Figure 5C). On the other hand, *parS*-specific variants (dark green box) have an invariable R at position 173, same as ParB orthologs (dashed dark green box) (Figure 5C), but position 201 can be small polar amino acids (C, S, or T, but G is most preferred). Notably, 17 amino acids (except the helix-breaking P or the negatively charged D and E) can occupy position 179, and any of the 20 amino acid is tolerable at position 184 (Figure 5C). Finally, dual-specificity variants (black box) tend to harbor sequence elements from both *parS*- and *NBS*-specific variants (Figure 5C).

### *NBS*-specific variants predominantly have lysine or arginine at positions 179 and 184

The proportion of *NBS*-specific variants with a K or R amino acid at position 179 is ∼58%, higher than a theoretical 10% value if K/R was chosen randomly (Figure 6A). The same proportion was seen for a K or R at position 184 (Figure 6A). This proportion increased to ∼91% for *NBS*-specific variants with either K or R at either position 179 or 184, and ∼19% for those with a K or R at both 179 and 184 (Figure 6A). The prevalence of positively charged residues, together with the structure of Noc (DBD)-*NBS*, supports our model that permissive mutations act by increasing protein-DNA binding affinity non-specifically via their interactions with a negatively charged phosphate backbone. We noted that K and R are not preferred more than expected from a random chance in *parS*-specific variants (Figure 6A). Our results suggest that the introduction of permissive substitutions is important to acquire a new specificity.

**Figure 6.**
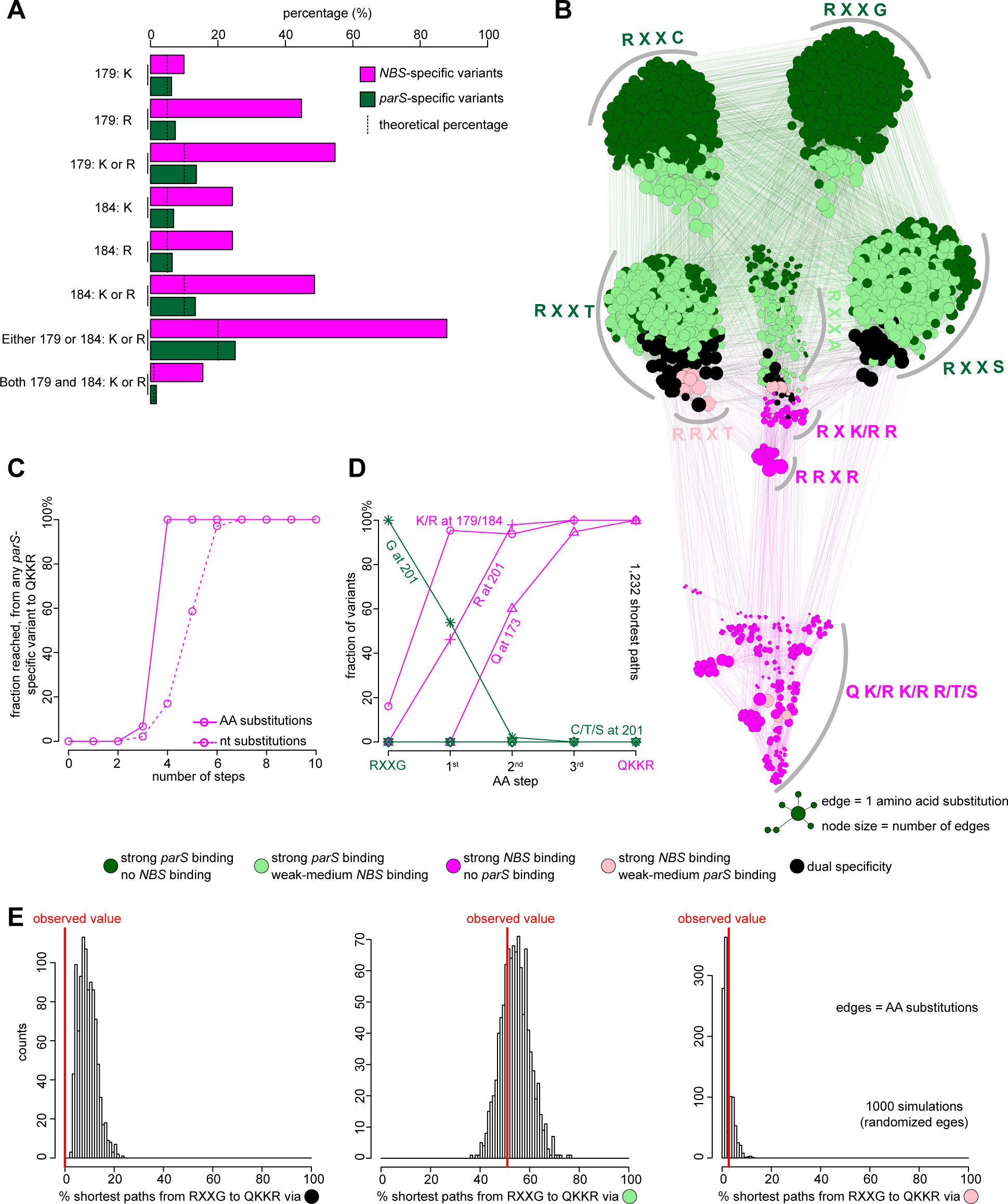
Deep mutational scanning experiments reveals the common properties of the mutational paths to a new DNA-binding specificity. **(A)** Fractions of arginine or lysine residues at position 179, 184, or both, in *parS*-specific (dark green) and *NBS*-specific (magenta) variants. The dotted lines indicate the expected percentage if arginine/lysine was chosen randomly from 20 amino acids. **(B)** A force-directed network graph connecting strong *parS*-binding variants to strong *NBS*-binding variants. Nodes represent individual variants, and edges represent single amino acid (AA) substitutions. Node sizes are proportional to their corresponding numbers of edges. Node colors correspond to different classes of variants. **(C)** Cumulative fraction of highly *parS*-specific variants that reached an *NBS*-specific QKKR variant in a given number of amino acid (solid line) or nucleotide (dotted line) substitutions (see also Figure S7A). **(D)** Fraction of intermediates on all shortest paths from highly *parS*-specific RXXG variants to the *NBS*-preferred QKKR that have permissive amino acids (K/R) at either position 179/184 or both, or have R at position 201, or Q at position 173, or C/T/S at position 201 after a given number of AA steps (see also Figure S7D). **(E)** Percentage of shortest paths that traversed black, light green, or pink variants to reach QKKR from any of the highly *parS*-specific RXXG variants (red lines). The result was compared to ones from 1,000 simulations where the edges were shuffled randomly while keeping the total number of nodes, edges, and graph density constant.

### Mutations were introduced in a defined order to reprogram specificity

We asked if there is an order of substitutions at positions 173, 179, 184, and 201 to create an *NBS*-specific variant. To answer this question, we first reconstructed all possible mutational paths to an *NBS* specificity. We created a force-directed graph that connects functional variants (nodes) together by lines (edges) if they are different by a single amino acid (AA) to visualize the connectivity of functional variants in sequence space (Figure 6B) (Podgornaia and Laub, 2015). The node size is proportional to its connectivity (number of edges), and node colors represent different classes of functional variants (Figure 6B). Similarly, we also generated a network graph in which edges represent variants that differ by a single nucleotide (nt) substitution (Figure S7A-B). Because not all amino acids can be converted to others by a mutation at a single base, a by-nt-substitution network might depict better how long (hard) or short (easy) the mutational paths that *parS*-specific variants might have taken to reprogram their specificity to *NBS*. At first glance, the network is composed of multiple clusters of densely interconnected nodes that share common features in the amino acid sequence (Figure 6B). Furthermore, there are multiple edges connecting *parS*-preferred variants (dark and light green nodes) to *NBS*-preferred variants (magenta and pink nodes) (Figure 6B). Supporting this observation, we found that it takes at most four AA (or seven nt) substitutions to convert any *parS*-specific variant to an *NBS*-specific QKKR (Figure 6C and Figure S7C). A small number of steps suggested that *NBS*-specific variants can be reached relatively easily from *parS*-specific variants. We focused on *parS*-specific start points RXXG for all analyses below because R173 and G201 are absolutely conserved in all extant ParB orthologs (Figure 5C). We found all the shortest paths (1,232 in total) that connect *parS*-specific RXXG variants (298 dark green nodes) to an *NBS*-specific QKKR, and quantified the fractions of intermediates in such paths that contain permissive or specificity-switching residues (Figure 6D). We discovered that permissive substitutions (K or R) at position 179 or 184 happened very early on along the mutational paths (∼95% after the first step, Figure 6D). The fraction of R201 increased more gradually after the introduction of permissive substitutions, and Q173 was introduced last (Figure 6D). The same order of substitutions was seen when we analyzed a by-nt-substitution network graph (Figure S7D). In sum, we conclude that the order of amino acid substitutions matters, and suggest that permissive mutations tend to happen before specificity-switching substitutions.

### Mutational paths that reprogram specificity did not travel across dual-specificity intermediates

We observed that the fraction of variants with C/T/S residues at position 201 did not increase beyond 0% in any step from RXXG variants to QKKR (Figure 6D and Figure S7D). Given that dual-specificity variants (black box, Figure 5C) mostly have T or S amino acid at position 201, it suggests that dual-specificity intermediates might have not been exploited to change specificity. Indeed, no shortest path connecting RXXG and QKKR traversed through any dual-specificity variant (black nodes) (Figure 6E and Figure S7E). This proportion is significantly smaller than would be expected by chance (estimated from 1,000 random networks where edges were shuffled randomly, Figure 6E and Figure S7E). In contrast, ∼51% and ∼3% of shortest paths from RXXG variants to QKKR contain light green and pink intermediates, respectively. The proportions of paths with light green or pink intermediates are similar to expected values from random chances (Figure 6E). The preference for traversing light green nodes, therefore, can be explained by the abundance of such variants in the observed graph (Figure 6B). Overall, our network analysis predicted that the *parS*-to-*NBS* reprogram did not exploit truly dual-specificity intermediates, and that those with a stricter specificity (light green or pink) were more commonly used.

## DISCUSSION

### Determinants of specificity and implications for understanding the evolution of protein-DNA interfaces

The *NBS* site differs from the *parS* site by only 2 bases (positions 1 and 6, Figure 1A) but Noc and ParB recognize and bind them with exquisite specificity. We provided evidence that mutations must have been introduced in a defined order to reprogram specificity. Permissive substitutions (K/R at positions 179/184) tend to appear first, presumably to prime *parS*-specific variants for a subsequent introduction of specificity-switching residues (R201 and Q173) which would have otherwise rendered proteins non-functional (Figure 7). Supporting the priming role of permissive amino acids, we noted that ∼28% of extant ParB already possess a lysine/arginine residue at position 184 (Figure 5C, a sequence logo in a dashed green box). An early introduction of permissive substitutions is likely to be a recurring principle of evolution. For example, a similar prerequisite for permissive mutations were observed in the evolution of influenza resistance to the antiviral drug oseltamivir [31]. Two permissive mutations were first acquired, allowing the virus to tolerate a subsequent occurrence of a H274Y mutation that weakened the binding of oseltamivir to the viral neuraminidase enzyme [31]. These permissive mutations improved the stability of neuraminidase before a structurally destabilizing H274Y substitution was introduced [31,32]. Similarly, a permissive mutation that is far away from the active site of an antibiotic-degrading β-lactamase (TEM1) has little effect on its enzymatic activity by itself but restored stability loss by a subsequent mutation that increased TEM1 activity against cephalosporin antibiotics [33]. In another case, 11 permissive mutations were required to evolve an ancestral steroid hormone receptor from preferring an estrogen response element (ERE) to a new DNA sequence (steroid response element or SRE) [34]. These 11 mutations were located outside of the DNA-recognition motif but non-specifically increased the affinity for both ERE and SRE, thereby licensing three additional substitutions to alter the specificity to SRE [34]. Additionally, it has been shown that an early introduction of 11 permissive substitutions dramatically increased the number of SRE-binding variants well beyond the historically observed variants [35]. In our work, at least when considering just four amino acid residues, a single introduction of a lysine, either at position 179 or 184, was sufficient to permit Q173 and R201 to recognize *NBS* specifically.

**Figure 7.**
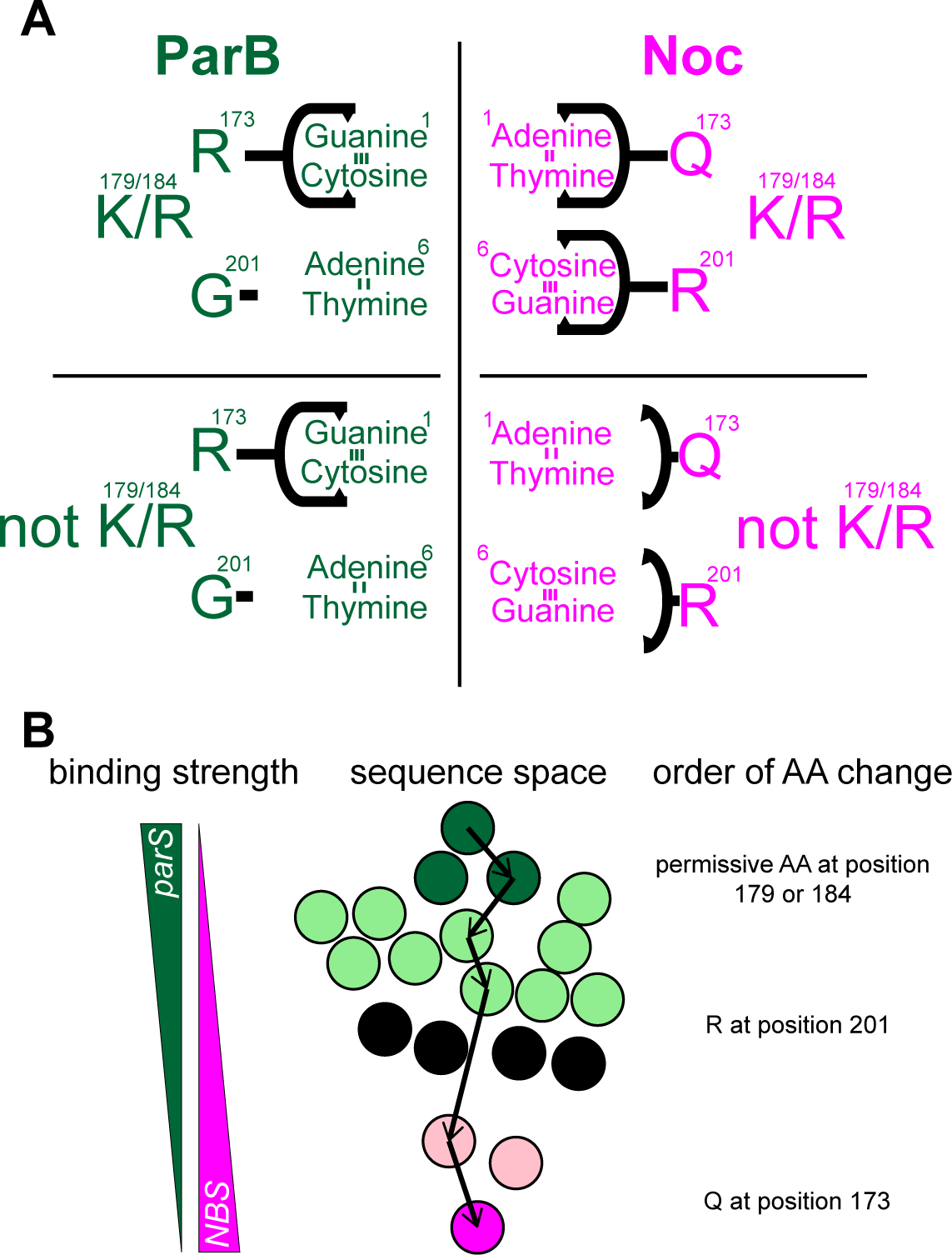
A model for the evolution of *NBS*-binding specificity. **(A)** Contributions of each specificity residue to enable a switch in binding specificity from *parS* to *NBS*. An R173Q substitution enabled interactions with Adenine 1:Thymine −1 (of *NBS*). A G201R substitution enabled interactions with Cytosine 6: Guanine −6 (of *NBS*). Q173 and R201 could only do so in the presence of permissive residues K at either 179, 184, or both. Without K179/184, Q173 and R201 were poised to interact with specific bases but could not, possibly because of insufficient affinity for DNA. **(B)** Analysis of mutational paths that traversed the network of functional variants showed that the order of introducing specificity-switching substitutions matters, and that the shortest paths to *NBS*-specific variants do not necessarily involve a dual-specificity nodes to evolve a new DNA-binding preference.

Deep mutational scanning in conjunction with network analysis is a powerful approach to reconstruct possible mutational paths that might have been taken to acquire a new function [30,35,36]. Network graph theory was applied to understand the constraints on the evolution of protein-protein interfaces between a histidine kinase and its response regulator partner, between toxin and antitoxin pairs of proteins, and most recently to reveal the alternative evolutionary histories of a steroid hormone receptor [30,35,36]. In our case study, network analysis suggested that mutational paths to a new specificity did not necessarily have to visit dual-specificity intermediates i.e. those that bind *parS* and *NBS* equally strongly (Figure 6E). Instead, mutational paths to an *NBS*-specific variant tend to be more switch-like, frequently visited dark green nodes (strong *parS* binding, no *NBS* binding) and light green nodes (strong *parS* binding, weak-to-medium *NBS* binding) (Figure 5C and Figure 6E). We reason that most black variants, albeit being dual specific, bind both *parS* and *NBS* at a slightly reduced affinity (compared to the wild-type *parS*-specific RTAG or *NBS*-specific QKKR variants, see the scatter plot on Figure 5C). This might have created an undesirable situation where dual-specificity intermediates neither could compete with the original copy of ParB to bind *parS* nor had high enough affinity themselves to bind *NBS* sites i.e. artificially made non-functional due to competition. A similar principle might also apply to other protein-DNA interactions throughout biology. For example, a reconstructed evolutionary history of a steroid hormone receptor indicated that an ancestral receptor (AncSR1) without permissive mutations must always pass through dual-specificity intermediates to acquire the present-day specificity. On the other hand, the presence of 11 permissive mutations (AncSR1+11P) eliminated the absolute requirement for these dual-specificity intermediates. More dramatically, it has been shown that a single substitution (i.e. a truly switch-like mechanism) was enough to reprogram the specificity of homologous repressor proteins (Arc and Mnt) in bacteriophage P22 [37]. Nevertheless, we noted that protein-protein interfaces, particularly in the case of paralogous toxin-antitoxin protein pairs, exploited extensively promiscuous intermediates to diversify and evolve instead. In the case of toxin-antitoxin systems, truly promiscuous intermediates might have been favored because many of them bound to and antagonized cognate and non-cognate toxins equally or even better than wild type [36]. It is likely that the topology of the available sequence space and the biology of each system collectively influence the paths to evolve a new biological innovation.

In sum, our work provides a molecular basis for how protein-DNA interaction specificity can change, with a focus on chromosome maintenance proteins ParB/Noc and the minimal set of four specificity residues at their protein-DNA interfaces. A small number of specificity residues enabled a systematic analysis of the protein-DNA interface and possible mutational paths that could have changed specificity. In this regard, our work might be useful in understanding the evolution of other classes of DNA-binding proteins. Nevertheless, evolution has most likely exploited more mutations and amino acid residues to fine-tune DNA-binding specificity than the core set of four residues in this work. Other compensatory mutations that alter the structural stability of proteins might also contribute and dictate the course of evolution to new biological functions [38–40]. An important challenge for future work is to study all contributing factors in a systematic manner to better understand the course of evolution to new biological innovations.

## ACKNOWLEDGEMENTS

This study was supported by the Royal Society University Research Fellowship (UF140053) and a BBSRC grant (BB/P018165/1) to T.B.K.L. A.S.B.J’s PhD studentship was funded by the Royal Society (RG150448), and N.T.T was funded by the BBSRC grant-in-add (BBS/E/J/000PR9791 to the John Innes Centre). Work in Dr. Agnes Noy lab was supported by EPSRC grant (EP/N027639/1). Computational time was secured via HECBioSim, supported by EPSRC grant (EP/R029407/1) and via Cambridge Tier-2 system (EPSRC Tier-2 capital grant EP/P020259/1). We acknowledge Tier 3 High Performance Computing facilities at York (Viking and YARCC clusters) for additional computational resources. We acknowledge Diamond Light Source for access to beamlines I03 and I04 under proposal MX18565 with support from the European Community’s Seventh Framework Program (FP7/2007–2013) under Grant Agreement 283570 (BioStruct-X). We also thank Rory Williams and Roan Hulks for help with early experiments

## AUTHOR CONTRIBUTIONS

Conceptualization: A.S.B.J, N.T.T, and T.B.K.L; Data acquisition: A.S.B.J, N.T.T, C.E.S, E.W.C, R.L, X.T, and T.B.K.L; Data analysis: A.S.B.J, N.T.T, C.E.S, E.W.C, R.L, A.N, D.M.L, and T.B.K.L. Writing: A.S.B.J, N.T.T, D.M.L, and T.B.K.L. Funding acquisition: A.N, D.M.L, and T.B.K.L.

## DECLARATION OF INTERESTS

The authors declare no competing interests.

## KEY RESOURCES TABLE

| REAGENT or RESOURCE | SOURCE | IDENTIFIER |
| --- | --- | --- |
| Bacterial and Virus Strains |  |  |
| Please refer to Table S1 |  | N/A |
| Chemicals, Peptides, and Recombinant Proteins |  |  |
| 3-AT | Abcam | Cat# ab146281 |
| Benzonase nuclease | Sigma-Aldrich | Cat# E1014 |
| cOmplete, Mini, EDTA-free Protease Inhibitor Cocktail | Roche Applied Science | Cat# 11836170001 |
| Critical Commercial Assays |  |  |
| Gibson Assembly Master Mix | NEB | Cat# E2611S |
| Gateway BP Clonase II enzyme mix | ThermoFisher | Cat# 11789020 |
| SYBR® Green JumpStart Taq ReadyMix | Sigma-Aldrich | Cat# S4438 |
| Dip-and-Read Streptavidin (SA) biosensors | Molecular Devices | Cat# 18-5019 |
| HisTrap High Performance column | GE Healthcare | Cat# GE17524801 |
| HiTrap Heparin High Performance column | GE Healthcare | Cat# GE17040601 |
| HiLoad 16/600 Superdex 75pg column | GE Healthcare | Cat# GE28989333 |
| HIS-Select Cobalt Affinity Gel | Sigma-Aldrich | Cat# H8162 |
| PD 10 Desalting Columns | Sigma-Aldrich | Cat# GE17085101 |
| EZview Red ANTI-FLAG® M2 Affinity Gel | Sigma-Aldrich | Cat# F2426 |
| QIAquick PCR Purification Kit | QIAGEN | Cat# 28104 |
| NEBNext Ultra™ DNA Library Prep Kit for Illumina | NEB | Cat# E7370S |
| Qubit™ dsDNA HS Assay Kit | ThermoFisher | Cat# Q32851 |
| Deposited Data |  |  |
| Crystal structures | This paper | PDB: 6S6H and PDB: 6Y93 |
| ChIP-seq data | This paper | GSE129285 (Gene Expression Omnibus) |
| B1H-seq data | This paper | GSE129285 (Gene Expression Omnibus) |
| Other data | This paper | Mendeley Data repository<br>DOI: 10.17632/8v45kvdtw5.1 |
| Recombinant DNA |  |  |
| Please refer to Table S2 |  | N/A |
| Oligonucleotides |  |  |
| Please refer to Table S3 |  | N/A |
| Software and Algorithms |  |  |
| BLItz Pro | Molecular Devices | Cat# 50-0156 |
| Gephi | The Open Graph Viz Platform | https://gephi.org/ |
| R | R Foundation for Statistical Computing | https://www.r-project.org/ |
| igraph | Igraph-The network analysis package | https://igraph.org/ |
| HHsuite | [41] | https://github.com/soedinglab/hh-suite |
| WebLogo 3.0 | [42] | http://weblogo.threeplusone.com/ |
| iTOL | [43] | https://itol.embl.de/ |
| CIPRES | [44] | http://www.phylo.org/ |
| Curves+ | [45] | http://curvesplus.bsc.es/analyse |
| Bowtie 1 | [46] | http://bowtie-bio.sourceforge.net/index.shtml |
| BEDTools | [47] | https://bedtools.readthedocs.io/en/latest/ |
| FASTX-Toolkit | Hannon Lab | http://hannonlab.cshl.edu/fastx_toolkit/ |
| DIALS | [48] | https://dials.github.io/ |
| AIMLESS | [49] | http://www.ccp4.ac.uk/html/aimless.html |
| CCP4i2 | [50] | http://www.ccp4.ac.uk/ |
| COOT | [51] | https://www2.mrc-lmb.cam.ac.uk/personal/pemsley/coot/ |
| PHASER | [52] | https://www.phenix-online.org/ |
| REFMAC5 | [53] | http://www.ccp4.ac.uk/html/refmac5/description.html |
| BUCCANEER | [54] | http://www.ccp4.ac.uk/html/cbuccaneer.html |
| MolProbity | [55] | http://molprobity.biochem.duke.edu/ |
| STARANISO | Global Phasing Limited | http://staraniso.globalphasing.org/cgi-bin/staraniso.cgi |
| jsPISA | [56] | http://www.ccp4.ac.uk/pisa/ |
| PyMOL | The PyMOL Molecular Graphics System | https://pymol.org/2/ |
| AMBER | AMBER Software, University of California, San Francisco | https://ambermd.org/ |

## MATERIALS & METHODS

*Escherichia coli* strains DH5α and Rosetta (DE3) were used as hosts for constructing plasmids, and overexpression of proteins, respectively (Table S1). *E. coli* USO *rpoZ*^*-*^ *hisB*^*-*^ *pyrF*^*-*^ was used as a host for B1H assay (Table S1).

### Growth conditions

*E. coli* was grown in LB. When appropriate, media were supplemented with antibiotics at the following concentrations (liquid/solid media for *E. coli* (μg/mL): carbenicillin (50/100), chloramphenicol (20/30), kanamycin (30/50), and apramycin (25/50).

### Plasmids and strains construction

All strains used are listed in Table S1. All plasmids and primers used in strain and plasmid construction are listed in Table S2 and S3.

#### pENTR::Noc/ParB

The coding sequences of ParB and Noc from various bacterial species (Figure 1B and Figure S1A) were chemically synthesized (gBlocks dsDNA fragments, IDT). The backbone of pENTR plasmid was amplified by PCR using primers pENTR_gibson_backbone_F and pENTR_gibson_backbone_R from the pENTR-D-TOPO cloning kit (Invitrogen). The resulting PCR product was subsequently treated with DpnI to remove methylated template DNA. The resulting PCR fragment was gel-purified and assembled with the gBlocks fragment using a 2x Gibson master mix (NEB). Gibson assembly was possible due to a 23 bp sequence shared between the PCR fragment and the gBlocks fragment. These 23 bp regions were incorporated during the synthesis of gBlocks fragments. The resulting plasmids were sequence verified by Sanger sequencing (Eurofins, Germany).

#### pUT18C-1xFLAG-DEST

The backbone of pUT18C was amplified using primers P1936 and P1937, and pUT18C [57] as template. The resulting PCR product was subsequently treated with DpnI to remove the methylated template DNA. The FLAG-*attR1*-*ccdB*-chloramphenicol^R^-*attR2* cassette was amplified using primers P1934 and P1935, and pML477 as template. The two PCR fragments were each gel-purified and assembled together using a 2x Gibson master mix (NEB). Gibson assembly was possible due to a 23 bp sequence shared between the two PCR fragments. These 23 bp regions were incorporated during the primer design to amplify the FLAG-*attR1*-*ccdB*-chloramphenicol^R^-*attR2* cassette. The resulting plasmid was sequence verified by Sanger sequencing (Eurofins, Germany).

#### pUT18C::1xFLAG-Noc/ParB

The *parB*/*noc* genes were recombined into a Gateway-compatible destination vector pUT18C-1xFLAG-DEST via a LR recombination reaction (Invitrogen). For LR recombination reactions: 1 µL of purified pENTR::*parB/noc* was incubated with 1 µL of the destination vector pUT18-1xFLAG-DEST, 1 µL of LR Clonase II master mix, and 2 µL of water in a total volume of 5 µL. The reaction was incubated for an hour at room temperature before being introduced to DH5α *E. coli* cells by heat-shock transformation. Cells were then plated out on LB agar + carbenicillin. Resulting colonies were restruck onto LB agar + carbenicillin and LB agar + kanamycin. Only colonies that survived on LB + carbenicillin plates were subsequently used for culturing and plasmid extraction.

#### pB1H2-w2::Caulobacter ParB (R104A + Q173K179K184R201) and pB1H2-w2::Caulobacter ParB (R104A + R173A179T184G201)

The coding sequence of *Caulobacter* ParB with the desired mutations was chemically synthesized (gBlocks dsDNA fragments, IDT). The pB1H2-w2 plasmid backbone was generated via a double digestion of pB1H2-w2::Prd plasmid [28] with KpnI and XbaI. The resulting backbone was subsequently gel-purified and assembled with the gBlocks fragments using a 2x Gibson mastermix (NEB). Gibson assembly was possible due to a 23 bp sequence shared between the KpnI-XbaI-cut pB1H2-w2 backbone and the gBlocks fragment. These 23 bp regions were incorporated during the synthesis of gBlocks fragments. The resulting plasmids were sequence verified by Sanger sequencing (Eurofins, Germany).

#### pB1H2-w5::Caulobacter ParB (R104A + Q173K179K184R201)

The same procedure as above was used to generate this plasmid, except that pB1H2-w5::Prd plasmid [28] was used.

#### pB1H2-w5L::Caulobacter ParB (R104A + Q173K179K184R201)

The same procedure as above was used to generate this plasmid, except that pB1H2-w5L::Prd plasmid [28] was used.

#### pU3H3::7/14/19/24bp-NBS

The pU3H3 backbone was generated via a double digestion of pU3H3::MCS plasmid [28] with XmaI and EcoRI. The backbone was subsequently gel-purified before being ligated with the DNA insert in the next step. The DNA insert containing *NBS* site with an appropriate spacer (7, 14, 19, or 24 bp) were generated by annealing complementary oligos together (Table S3). The DNA inserts were subsequently 5’ phosphorylated using T4 PNK (NEB), and ligated to the XmaI-EcoRI-cut pU3H3 backbone using T4 DNA ligase (NEB). The resulting plasmids were sequence verified by Sanger sequencing (Eurofins, Germany).

#### pU3H3::19bp-parS

The same procedure as above was used to generate the plasmid, except that primers *parS*_anneal_19bp_spacer_F and *parS*_anneal_19bp_spacer_R were used.

#### pET21b::ParB (variants)-His_6_

All sequences of ParB variants were designed in VectorNTI (ThermoFisher) and chemically synthesized as gBlocks dsDNA fragments (IDT). Individual gBlocks fragment and a NdeI-HindIII-digested pET21b backbone were asssembled using a 2x Gibson master mix (NEB). Gibson assembly was possible due to a 23-bp sequence shared between the NdeI-HindIII-cut pET21b backbone and the gBlocks fragment. These 23-bp regions were incorporated during the synthesis of gBlocks fragments. The resulting plasmids were sequence verified by Sanger sequencing (Eurofins, Germany).

#### Strains AB1157 ybbD::parS::markerless ygcE::NBS::markerless

We use Lambda Red to insert a cassette consisting of a *parS* site and an apramycin antibiotic resistance gene *aac(3)IV* at the *ybbD* locus on the *E. coli* AB1157 chromosome. The *parS*-FRT-apramycin^R^-FRT cassette was amplified by PCR using primers 1940 and 1941, and pIJ773 (a gift from Keith Chater) as template. These forward and reverse primers also carry a 39 bp homology to the left or the right of the insertion point at the *ybbD* locus. The resulting PCR products were gel-extracted and electroporated into an arabinose-induced *E. coli* AB1157/pKD46 cells. Colonies that formed on LB + apramycin was restruck on LB + apramycin and incubated at 42°C to cure the cells of pKD46 plasmid. Finally, the correct insertion of the *parS*-apramycin^R^ cassette was verified by PCR and Sanger sequencing. To remove the FRT-apramycin^R^-FRT region while leaving the *parS* site intact, a temperature sensitive FLP recombination plasmid pBT340 (a gift from Keith Chater) was subsequently introduced. To introduce the *NBS* site at the *ygcE* locus on the chromosome of *E. coli* AB1157 *ybbD*::*parS*::markerless, we employed the same procedure, except that the *NBS*-FRT-Apramycin^R^-FRT cassette was amplified by PCR using primer 3139 and 3140 instead.

## METHOD DETAILS

### Identification and alignment of ParB and Noc sequences

The sequences used for generating sequence conservation logos were retrieved and aligned using HHblits (-n 4 -e 1E-10 -maxfilt inf -neffmax 20 -nodiff -realign_max inf) and HHfilter (-id 100 –cov 75) in the HHsuite [41], using *Caulobacter crescentus* ParB protein and *Bacillus subtilis* Noc protein sequences as queries. This procedure resulted in 1800 homologous ParB sequences and 361 homologous Noc sequences. The sequence conservation logos were generated by WebLogo 3.0 [42], using ParB/Noc sequence alignments as input.

### Phylogenetic analysis of ParB and Noc protein sequences

Amino acid sequences of ParB and Noc from 21 selected bacterial species were retrieved by BLASTP and used to generate a phylogenetic tree (Figure 1B). Phylogenetic analyses were carried out using MUSCLE [58] and RAxML [59], which were used through the CIPRES science gateway [44], and the trees were visualized using iTOL [43]. Amino acid sequences were aligned using MUSCLE with the following parameters: muscle -in infile.fasta -seqtype auto -maxiters 16 - maxmb 30000000 -log logfile.txt -weight1 clustalw -cluster1 upgmb -sueff 0.1 -root1 pseudo - maxtrees 1 -weight2 clustalw -cluster2 upgmb -sueff 0.1 -root2 pseudo -objscore sp -noanchors - phyiout outputi.phy

The resulting PHYLIP interleaved output file was then used to generate a maximum likelihood phylogenetic tree using RAxML-HPC BlackBox. The program was configured to perform rapid bootstrapping, followed by a maximum likelihood search to identify the best tree, with the following input parameters: raxmlHPC-HYBRID_8.2.10_comet -s infile.phy -N autoMRE -n result -f a -p 12345 -x 12345 -m PROTCATJTT

### Protein overexpression and purification

The DNA-binding domain (DBD) of *Caulobacter* ParB (residues 126-243) was expressed and purified as follows. Plasmid pET21b::*Caulobacter crescentus*-ParB-(His)_6_ (residue 126-243) was introduced into *E. coli* Rosetta (DE3) competent cells (Merck) by heat-shock transformation. 10 mL overnight culture was used to inoculate 4 L LB medium + carbenicillin + chloramphenicol. Cells were grown at 37°C with shaking at 210 rpm to an OD_600_ of ∼0.4. The culture was then left to cool to 28°C before isopropyl-β-D-thiogalactopyranoside (IPTG) was added at a final concentration of 1.0 mM. The culture was left shaking for an additional 3 hours at 28°C before cells were harvested by centrifugation.

Pelleted cells were resuspended in a buffer containing 100 mM Tris-HCl pH 8.0, 300 mM NaCl, 10 mM Imidazole, 5% (v/v) glycerol, 1 µL of Benzonase nuclease (Sigma Aldrich), 1 mg of lysozyme (Sigma Aldrich), and an EDTA-free protease inhibitor tablet (Roche). The pelleted cells were then lyzed by sonification (10 cycles of 15 s with 10 s resting on ice in between each cycle). The cell debris was removed though centrifugation at 28,000 g for 30 min and the supernatant was filtered through a 0.45 µm sterile filter. The protein was then loaded into a 1-mL HiTrap column (GE Healthcare) that had been equilibrated with buffer A [100 mM Tris-HCl pH8.0, 300 mM NaCl, 10 mM Imidazole, and 5% glycerol]. Protein was eluted from the column using an increasing (10 mM to 500 mM) imidazole gradient in the same buffer. ParB (DBD)-containing fractions were pooled and diluted to a conductivity of 16 mS/cm before being loaded onto a Heparin HP column (GE Healthcare) that had been equilibrated with 100 mM Tris-HCl pH 8.0, 25 mM NaCl, and 5% glycerol. Protein was eluted from the Heparin column using an increasing (25 mM to 1 M NaCl) salt gradient in the same buffer. ParB (DBD) fractions were pooled and analyzed for purity by SDS-PAGE. Glycerol was then added to ParB fractions to a final volume of 10%, followed by 10 mM EDTA and 1 mM DDT. The purified ParB (DBD) was subsequently aliquoted, snap frozen in liquid nitrogen, and stored at −80°C. ParB (DBD) that was used for X-ray crystallography was further polished via a gel-filtration column. To do so, purified ParB (DBD) was concentrated by centrifugation in an Amicon Ultra-15 3-kDa cut-off spin filters (Merck) before being loaded into a Superdex 75 gel filtration column (GE Healthcare). The gel filtration column was pre-equilibrated with 10 mM Tris-HCl pH 8.0 and 250 mM NaCl. ParB (DBD) fractions was then pooled and analyzed for purity by SDS-PAGE.

The DNA-binding domain (DBD) of *Bacillus subtillis* Noc-(His)_6_ (residue 111-242) was purified using the same 3-column procedure as above.

All other ParB/Noc variants were purified using HIS-Select^®^ Cobalt gravity flow columns as follows. Plasmid pET21b::*parB/noc* variants were introduced individually into *E. coli* Rosetta (DE3) competent cells (Merck) by heat-shock transformation. 10 mL overnight culture was used to inoculate 1 L LB medium + carbenicillin + chloramphenicol. Cells were grown at 37°C with shaking at 210 rpm to an OD_600_ of ∼0.4. The culture was then left to cool to 28°C before IPTG was added to a final concentration of 0.5 mM. The culture was left shaking for an additional 3 hours at 30°C before cells were harvested by centrifugation. Pelleted cells were resuspended in 25 ml of buffer A [100 mM Tris-HCl pH 8.0, 300 mM NaCl, 10 mM Imidazole, 5% (v/v) glycerol] containing 1 mg lysozyme (Sigma Aldrich), and an EDTA-free protease inhibitor tablet (Roche). The pelleted cells were then lyzed by sonification. The cell debris was removed though centrifugation at 28,000 g for 30 min and the supernatant was transferred to a gravity flow column containing 2 mL of HIS-Select^®^ Cobalt Affinity Gel (Sigma Aldrich) that was pre-equilibrated with 40 mL of buffer A. The column was rotated at 4°C for 1 hour to allow for binding to His-tagged proteins to the resin. After the binding step, unbound proteins were washed off using 60 mL of buffer A. Proteins were eluted using 2.7 mL of buffer B [100 mM Tris-HCl pH 8.0, 300 mM NaCl, 500 mM Imidazole, 5% (v/v) glycerol]. The purified protein was desalted using a PD-10 column (GE Healthcare), concentrated using an Amicon Ultra-4 10 kDa cut-off spin column (Merck), and stored at −80°C in a storage buffer [100 mM Tris-HCl pH 8.0, 300 mM NaCl, and 10% (v/v) glycerol].

### Selection of *parS* and *NBS* site

For all experiments described in this work, we employed a consensus *parS* site (TGTTTCACGTGAAACA) and consensus *NBS* site (TATTTCCC-GGGAAATA) i.e. the idealized sequence that represents the predominant base at each position. The full position weight matrix (PWM) logos for *parS* and *NBS* sites have been described previously [6,25].

### Reconstitution of *parS* DNA for X-ray crystallography

A 20-bp palindromic DNA fragment (5’-GAT<u>GTTTCACGTGAAAC</u>ATC-3’) (3.6 mM in buffer that contains 10 mM Tris-HCl pH 8.0 and 250 mM NaCl) was heated to 95°C for 5 min before being left to cool at room temperature overnight to form a double stranded *parS* DNA (final concentration: 1.8 mM). The 14-bp *parS* site sequences are underlined.

### Reconstitution of *NBS* DNA for X-ray crystallography

A 22-bp DNA fragment (5’-GGAT<u>ATTTCCCGGGAAAT</u>ATCC-3’) (3.6 mM in buffer that contains 10 mM Tris-HCl pH 8.0 and 250 mM NaCl) was heated to 95°C for 5 min before being left to cool at room temperature overnight to form a double stranded *NBS* DNA (final concentration: 1.8 mM). The 14-bp *NBS* site sequences are underlined.

### Protein crystallization, structure determination, and refinement

Crystallization screens were set up in sitting-drop vapour diffusion format in MRC2 96-well crystallization plates (Swissci) using either an OryxNano or an Oryx8 robot (Douglas Instruments) with drops comprised of 0.3 μL precipitant solution and 0.3 µL of protein-DNA complex, and incubated at 293 K. After optimisation of initial hits, suitable crystals were cryoprotected with 20% (v/v) glycerol and mounted in Litholoops (Molecular Dimensions) before flash-cooling by plunging into liquid nitrogen. X-ray data were recorded on either beamline I04 or I03 at the Diamond Light Source (Oxfordshire, UK) using either a Pilatus 6M-F or an Eiger2 XE 16M hybrid photon counting detector (Dectris), respectively, with crystals maintained at 100 K by a Cryojet cryocooler (Oxford Instruments). Diffraction data were integrated and scaled using DIALS [48] via the XIA2 expert system [60] then merged using AIMLESS [49]. The Noc (DBD)-*NBS* data set was further subjected to anisotropic correction using the STARANISO server as detailed below. Data collection statistics are summarized in Supplementary Table S4. The majority of the downstream analysis was performed through the CCP4i2 graphical user interface [50].

#### DNA-binding domain (DBD) ParB in complex with 20-bp parS

For crystallization, His-tagged DBD ParB (10 mg/mL) was mixed with a 20-bp *parS* site at a molar ratio of 1:1.2 (protein:DNA) in the elution buffer (10 mM Tris-HCl pH 8.0, 250 mM NaCl). The DBD ParB-*parS* complex crystals grew in a solution containing 19% (w/v) PEG3350 and 49 mM lithium citrate.

The ParB (DBD)-*parS* complex crystallized in space group *C*2 with approximate cell parameters of *a* = 122.1, *b* = 40.7, *c* = 94.0 Å and *β* = 121.4° (Table S4). Analysis of the likely composition of the asymmetric unit (ASU) suggested that it would contain two copies of the ParB (DBD) bound to a single DNA duplex, giving an estimated solvent content of ∼49%. A molecular replacement template covering the DBD was generated by manually editing the protein component of the structure of the Spo0J-*parS* complex from *Helicobacter pylori* [61] (PDB accession code 4UMK; 42% identity over 75% of the sequence) and truncating all side-chains to Cβ atoms. For the DNA component, an ideal B-form DNA duplex was generated in COOT [51] from the 20-bp palindromic sequence of *parS*. PHASER [52] was used to place the DNA duplex, followed by two copies of the DBD template into the ASU. The placement of the DNA-binding domains with respect to the DNA duplex was analogous to that seen in the *Helicobacter* Spo0J-*parS* [61], and an analysis of crystal contacts revealed that the DNA formed a pseudo-continuous filament spanning the crystal due to base-pair stacking between adjacent DNA fragments. After restrained refinement in REFMAC5 [53] at 2.4 Å resolution, the protein component of the model was completely rebuilt using BUCCANEER [54]. The model was finalized after several iterations of manual editing in COOT and further refinement in REFMAC5 incorporating TLS restraints. The model statistics are reported in Table S4.

#### DNA-binding domain (DBD) Noc in complex with 22-bp NBS

Crystallization screens were set up in sitting-drop vapor diffusion format in MRC2 96-well crystallization plates with drops comprised of 0.3 µL precipitant solution and 0.3 µL of protein-DNA complex, and incubated at 293 K. Noc (DBD)-His_6_ (10 mg/mL) was mixed with a 22-bp *NBS* duplex at a molar ratio of 1:1.2 protein:DNA in buffer containing 10 mM Tris-HCl pH 8.0 and 250 mM NaCl. The Noc (DBD)-*NBS* crystals grew in a solution containing 20% (w/v) PEG 3350 and 200 mM di-potassium phosphate.

The Noc (DBD)-*NBS* complex crystallized in space group *C*2 with approximate cell parameters of *a* = 134.1, *b* = 60.6, *c* = 81.0 Å and *β* = 116.9°. The data were collected in two 360° sweeps separated by a χ offset of 20°. Data reduction in AIMLESS indicated that the diffraction was highly anisotropic, and thus before using the dataset, it was corrected using STARANISO with a local mean *I/σ(I)* threshold of 1.2, giving maximum and minimum anisotropic resolution cut-offs of 2.23 and 4.02 Å, respectively (Table S4). Analysis of the likely composition of the asymmetric unit (ASU) suggested that it would contain two copies of the Noc (DBD) bound to a single DNA duplex, giving an estimated solvent content of ∼69%. A molecular replacement template covering the DBD was generated from the ParB DBD structure above using SCULPTOR (41% identity overall) [62]. For the DNA component, an ideal B-form DNA duplex was generated from the 22-bp palindromic sequence of *NBS*. PHASER was used to place the DNA duplex, followed by two copies of the DBD template into the ASU. This generated a complex that was consistent with that of ParB (DBD)-*parS* determined above, again with the DNA forming a pseudo-continuous filament spanning the crystal due to base-pair stacking between adjacent DNA fragments. After restrained refinement in REFMAC5 at 2.23 Å resolution, the protein component of the model was completely rebuilt using BUCCANEER [54]. The model was finalized after several iterations of manual editing in COOT and further refinement in REFMAC5 incorporating TLS restraints. To avoid model bias resulting from the feature of REFMAC5 to approximate missing reflections within the spherical resolution cut-off to their calculated values, these filled-in reflections were removed prior to map inspection. Subsequently, the map connectivity was improved by applying a blurring factor of 60 Å^2^. The model statistics are reported in Table S4.

### Identification of protein-DNA contacts and analysis of DNA shapes

Protein-DNA contacts were identified using the jsPISA webserver [56]. Superpositions of structures were performed using the *align/cealign* function in PyMOL. DNA shape parameters were determined from the structures using Curves+ [45].

### Molecular dynamics simulations

We performed simulations of Noc (DBD)-*NBS* complex using its crystallographic structure as initial coordinates. Virginia Tech H++ web server [63] was used for ensuring the correct protonated state of proteins at pH 7.0. Forcefields ff14SB [64] and parmbsc1 [65] were employed for describing protein and DNA, respectively. The system was solvated in a TIP3P octahedral periodic box [66] with a 12 Å buffer and 100 mM of NaCl ions [67]. Minimization and equilibration were performed following a standard protocol [68] at constant temperature (300 K) and pressure (1 atm). The structures were simulated for 200 ns with an integration time step of 2 fs. SHAKE method [69] was used to constrain hydrogen bonds, alongside periodic boundary conditions and Particle-Mesh-Ewald algorithm [70]. These simulations were performed with CUDA implementation of AMBER 18’s PMEMD module. After discarding the first 10 ns, trajectory was analyzed using cpptraj [71] for describing the nature of protein:DNA interactions. Hydrogen bonds were determined using a distance cutoff of 3.5 Å between donor and acceptor atoms and an angle cutoff of 120°. Salt bridges were also established with a distance cutoff of 3.5 Å for a direct ion-pair contact between heavy atoms of charged groups and an increased cutoff of 6.0 Å for a solvent-separated ion-pair [72].

### Measure protein-DNA binding affinity by bio-layer interferometry (BLI)

Bio-layer interferometry experiments were conducted using a BLItz system equipped with Dip-and-Read Streptavidin (SA) Biosensors (ForteBio). BLItz monitors wavelength shifts (response, unit: nm) resulting from changes in the optical thickness of the sensor surface during association or dissociation of the analyte over time to obtain kinetics data i.e. k_off_ and k_on_ of interactions. The streptavidin biosensor (ForteBio) was hydrated in a binding buffer [100 mM Tris-HCl pH 7.4, 150 mM NaCl, 1 mM EDTA, and 0.005% Tween 20] for at least 10 min before each experiment. Biotinylated dsDNA was immobilized onto the surface of the SA biosensor through a cycle of Baseline (30 s), Association (120 s), and Dissociation (120 s). Briefly, the tip of the biosensor was dipped into a low salt buffer for 30 s to establish the baseline, then to 1 μM biotinylated dsDNA for 120 s, and finally to a low salt binding buffer for 120 s to allow for dissociation. Biotinylated dsDNA harboring *parS, NBS*, or variant of such sites were prepared by annealing a 24-bp biotinylated oligo with its unmodified complementary strand in an annealing buffer [1 mM Tris-HCl pH 8.0 and 5 mM NaCl]. The oligos mixture was heated to 98°C for 2 min and allowed to cool down to room temperature overnight.

After immobilizing DNA on the sensor, we first screened for protein-DNA interactions using a high protein concentration (1000 nM dimer concentration) (282 unique protein-DNA pairs in total, triplicated screens). A protein-DNA pair was regarded as not interacting if no/very weak BLI response above background was observed at this concentration, hence K_D_ was not determined. For other protein-DNA pairs where we observed BLI responses at 1000 nM, experiments were extended to include a range of protein concentrations. The concentration used were typically 0, 31, 62, 125, 250, 500, and 1000 nM. For weaker protein-DNA pairs, higher concentrations such as 2000 and 4000 nM were also employed. At the end of each protein binding step, the sensor was transferred into a protein-free binding buffer to follow the dissociation kinetics for 120 s. The sensor could be recycled by dipping in a high-salt buffer [100 mM Tris-HCl pH 7.4, 1000 mM NaCl, 1 mM EDTA, and 0.005% Tween 20] for at least 1 min to remove bound proteins.

For every protein-DNA pair, we first measured the kinetics (i.e. response vs. time) at 1000 nM in triplicate, using three independent protein aliquots. The kinetic profiles were deemed reproducible, with deviations less than 10%. Then, for Figure 2D and Figure 3C, we measured the kinetics once for each concentration (0, 31, 62, 125, 250, and 500 nM). Kinetics data were fitted locally, using an 1:1 binding model, for each protein concentration using BLItz Pro software (ForteBio) to determine k_off_, k_on_, and K_D_ (a ratio of k_off_/k_on_). The χ^2^ and R^2^ values were calculated, a local fitting was judged to be good if χ^2^<3 and R^2^>0.9. For a poor local fitting (i.e. χ^2^>3 and R^2^<0.9), typically because of a low BLI response at a low protein concentration, this datapoint was omitted from K_D_ calculation (BLI data analysis manual-ForteBio). Each calculated K_D_ at each concentration is considered as an independent determination of such value, hence we averaged to obtain mean K_D_ and standard deviation (S.D) for each protein-DNA pair. For Figure 3A, we measured the kinetics in triplicate for every concentration in the range.

### Clustering of trajectory-scanning mutagenesis data

K_D_ of interactions between ParB (WT)/PtoN variants and each of the 16 DNA-binding sites were presented as a two-dimensional heatmap using the *heatmap* function in R. Euclidean distances were measured to obtain a distance matrix, and a complete agglomeration method, implemented within the *heatmap* function, was used for clustering.

### Chromatin immunoprecipitation with qPCR or deep sequencing

For *E. coli* ChIP-seq, cells harboring pUT18C-1xFLAG-ParB/Noc were grown in LB (25 mL) at 28°C to mid exponential phase (OD_600_ ∼0.4) before 1 mM IPTG was added for 1-3 hours. The induction time (either 1, 2 or 3 hours) was chosen so that all ParB/Noc variants were produced to a similar level as judged by an α-FLAG Western blot. Subsequently, formaldehyde is added to a final concentration of 1% to fix the cells.

Fixed cells were incubated at room temperature for 30 min, then quenched with 0.125 M glycine for 15 min at room temperature. Cells were washed three times with 1x PBS (pH 7.4) and resuspended in 1 mL of buffer 1 [20 mM K-HEPES pH 7.9, 50 mM KCl, 10% Glycerol and Roche EDTA-free protease inhibitors]. Subsequently, the cell suspension was sonicated on ice using a probe-type sonicator (8 cycles, 15 s on 15s off, at setting 8) to shear the chromatin to below 1 kb, and the cell debris was cleared by centrifugation (20 min at 13,000 rpm at 4°C).

The supernatant was then transferred to a new 1.5 mL tube and the buffer conditions were adjusted to 10 mM Tris-HCl pH 8, 150 mM NaCl and 0.1% NP-40. Fifty microliters of the supernatant were transferred to a separate tube for control (the INPUT fraction) and stored at - 20°C. In parallel, antibodies-coupled beads were washed off storage buffers before adding to the above supernatant. We employed α-FLAG antibodies coupled to agarose beads (Sigma Aldrich) for ChIP-seq of FLAG-ParB/Noc. Briefly, 100 μL of beads was washed off storage buffer by repeated centrifugation and resuspension in IPP150 buffer [10 mM Tris-HCl pH 8, 150 mM NaCl and 0.1% NP-40]. Beads were then introduced to the cleared supernatant and incubated with gentle shaking at 4°C overnight. In the next day, beads were then washed five times at 4°C for 2 min each with 1 mL of IPP150 buffer, then twice at 4°C for 2 min each in 1x TE buffer [10 mM Tris-HCl pH 8 and 1 mM EDTA]. Protein-DNA complexes were then eluted twice from the beads by incubating the beads first with 150 μL of the elution buffer [50 mM Tris-HCl pH 8.0, 10 mM EDTA, and 1% SDS] at 65°C for 15 min, then with 100 μL of 1x TE buffer + 1% SDS for another 15 min at 65°C. The supernatant (the ChIP fraction) was then separated from the beads and further incubated at 65°C overnight to completely reverse crosslink. The INPUT fraction was also de-crosslinked by incubation with 200 μL of 1x TE buffer + 1% SDS at 65°C overnight. DNA from the ChIP and INPUT fraction were then purified using the PCR purification kit (Qiagen) according to the manufacturer’s instruction, then eluted out in 50 µL of EB buffer (Qiagen). The purified DNA was then used directly for qPCR or being constructed into library suitable for Illumina sequencing using the NEXT Ultra library preparation kit (NEB). ChIP libraries were sequenced on the Illumina HiSeq 2500 at the Tufts University Genomics facility. For the list of ChIP-seq datasets in this study, see Table S5.

### Generation and analysis of ChIP-seq profiles

For analysis of ChIP-seq data, Hiseq 2500 Illumina short reads (50 bp) were mapped back to the *Escherichia coli* MG1655 reference genome using Bowtie1 [46] and the following command: bowtie -m 1 -n 1 --best --strata -p 4 --chunkmbs 512 MG1655*-*bowtie --sam *.fastq > output.sam

Subsequently, the sequencing coverage at each nucleotide position was computed using BEDTools [47] using the following command: bedtools genomecov -d -ibam output.sorted.bam -g Ecoli_MG1655.fna > coverage_output.txt

ChIP-seq profiles were plotted with the x-axis representing genomic positions and the y-axis is the number of reads per base pair per million mapped reads (RPBPM) using custom R scripts. To calculate the enrichment of reads at the *parS* or *NBS* site (Figure S1B), we summed the RPBPM values for a 100-bp window surrounding the *parS* or *NBS* site.

### Bacterial one-hybrid assay coupled with deep sequencing (B1H-seq)

#### Optimization of bacterial one-hybrid assays

Bacterial one-hybrid (B1H) assays were performed as described previously [28]. Recipes for the minimal medium for B1H selection were described in detail previously [28]. Several parameters (promoter strength, spacers between the core −10 −35 promoter and the *NBS/parS* site, and IPTG concentration) were empirically optimized for experiments described in this work (Figure S4). We found that induction of *ω-parB*\* from a weak *lacUV5mut* promoter, using 0.1 mM IPTG, minimizes toxicity to the cells. Also, a 19-bp spacer between the core −10 −35 promoter and the *parS/NBS* site is optimal for the induction of HIS3 URA3 but does not auto-induce these genes (Figure S4). Therefore, we employed *pU3H3::19bp-parS* and *pU3H3::19bp-NBS* plasmids for all subsequent B1H selection.

#### Construction of combinatorial plasmid libraries

To construct combinatorial mutagenesis libraries where codons for Q173, K179, K184, and R201 were replaced with NNS (N = A/T/G/C, S=G/C), we employed round-the-horn PCR using oligos For_B_NNS_HTH, Rev_B_NNS_HTH, and pB1H2-P_*lacUV5mut*_*-Caulobacter* ParB (R104A + Q173K179K184R201) plasmid as template. Briefly, desalted oligos were reconstituted in 1x T4 ligase buffer, and 5’ phosphorylated using T4 PNK enzyme (NEB). Thirty 50µL PCR reactions were performed before DpnI was added and incubated overnight at 37°C to remove the methylated template. Next, PCR product (∼4.5 kb) was gel-purified and re-circularized overnight using T4 DNA ligase (NEB). The product was then ethanol precipitated to remove salts, and the DNA pellet was resuspended in 50 µL of water before being introduced into electrocompetent *E. coli* DH5α cells. Around 15 million carbenicillin-resistant *E. coli* colonies were collected, pooled together, and have their plasmid extracted (Qiagen MiniPrep kit). The whole procedure was repeated two more times, and on different days, to obtain three independent combinatorial libraries. Libraries from ∼15 million individual colonies ensure that at least 99% completeness is achieved [73].

#### Selection of ParB variants that bind to NBS or parS

The selection strain TLE3001 (USO *rpoZ- hisB- pyrF-*) harboring either *pU3H3::19bp-NBS* or *pU3H3::19bp-parS* plasmid was made electrocompetent. Next, approximately 2 µg of the combinatorial plasmid library were electroporated into 100 µL of the selection strain. The procedure was repeated for four more times, and electroporated cells were recovered in 10 mL of LB for an hour at 37°C. Subsequently, cells were washed off rich LB medium and resuspended in 5ml of 1x M9 liquid. Cells were then plated out on ten 150 mm Petri plates containing M9-minus-histidine medium supplemented with 0.1 mM IPTG, 5 mM 3-AT (a competitive inhibitor of HIS3, to increase the stringency of the selection), and appropriate antibiotics. Plates were incubated at 37°C for 48 hours before cells were scrapped off the agar surface, pooled together, and had their plasmids extracted (Qiagen Miniprep kit).

#### Construction of deep sequencing libraries

Illumina Truseq-compatible libraries were constructed from pre- and post-selection plasmid libraries via two rounds of PCR.

##### PCR round 1

Primer 4nns_R (10 µM): 2.5 µL

Mixture in equimolar amount of primers 4nns_offset_0_F; 4nns_offset_1_F; 4nns_offset_2_F; 4nns_offset_3_F; 4nns_offset_4_F (10 µM): 2.5 µL. A mixture of forward primers were used to stagger reads across the amplicon to improve the distribution of base calls at each position during the initial rounds of Illumina sequencing.

dNTP (10mM): 1 µL DMSO: 1.5 µL

Plasmid template (pre- or post-selection): 1 µL of 500 ng/µL Phusion polymerase: 0.5 µL

5x HF buffer: 10 µL

Water: 31 µL

PCR program: 98°C for 30 s, (98°C for 10 s, 56°C for 20 s, 72°C for 10 s) x 20 cycles, 72°C for 5 min.

PCR products were gel-purified, quantified by Qubit hsDNA quantification kit (ThermoFisher), and used as template in the second PCR.

##### PCR round 2

NEBNext Index primer (NEB): 2.5 µL

NEBNext universal primer (NEB): 2.5 µL

dNTP (10mM): 1 µL

DMSO: 1.5 µL

Template: 5 µL of gel-purified DNA from PCR round 1

5x HF buffer: 10 µL

Phusion polymerase: 0.5 µL

Water: 27 µL

PCR program: 98°C for 30s, (98°C for 10s, 54°C for 20 s, 72°C for 10 s) x 12 cycles, 72°C for 5 min.

PCR products were gel-purified, quantified by Qubit hsDNA quantification kit (ThermoFisher), and were sequenced on the Illumina HiSeq 2500 (single-end, 150-bp read length, 15% spike-in phiX DNA) at the Tufts University Genomics facility. For the list of B1H-seq datasets in this study, see Table S5.

### Analysis of data from deep mutational scanning experiments

#### Processing deep sequencing reads

We used fastx_trimmer script from the FASTX-Toolkit to remove nucleotides 0 to 20 and 114 to 150 from our reads using the following command: fastx_trimmer -f 20 -l 114 -Q33 -i TLE4_S4_R1_001.fastq -o TLE4_trimmed.fastq. Subsequently, we discarded sequence reads with an average Phred score <28, using the fastq_quality_filter script in the FASTX-Toolkit: fastq_quality_filter -v -Q33 -q 28 -p 100 -i TLE4_trimmed -o TLE4_trimmed_filtered.fastq. Reads were further filtered for the exact match to the following sequence:[ATGC][ATGC][GC]tctcacgtagcgaat[ATGC][ATGC][GC]atgcgtcttctt[ATGC][ATGC][GC]ttg ccggacgaggtacagtcctatcttgtgagtggagagctgacagcg[ATGC][ATGC][GC]. Corresponding codons (bases 1-3, 19-21, 34-36, 85-87) for the four specificity residues were extracted from the above 87-bp nucleotide sequence, and subsequently translated to amino acid sequence, following the standard genetic code. Variants with stop codon (TAG) were removed and were not considered in subsequent steps. Because of a high reproducibility among replicates (Figure S5C), we pooled reads from three replicates together (Figure S5A). We counted the number of occurrences (counts) for each unique variant, and removed variants with less than 10 reads (Figure S5B). Greater than 94% of all 160,000 predicted variants were represented by at least 10 reads. The variant counts for pre-selection and post-selection (for *parS*- or *NBS*-binding) libraries were used in the following steps to estimate the fitness score of each variant.

#### Calculation of fitness scores

We calculated the fitness of each variant (*f*_*parS*_ *and f*_*NBS*_), in comparison to WT variants (RTAG or QKKR), as described previously [36,74].

**f**_*parS, raw*_ *= log*_*10*_*(N variant, parS post-selection library / N wt, parS post-selection library) - log*_*10*_*(N variant, pre-selection library / N wt, pre-selection library)*

N variant, *parS* post-selection library = counts of each variant in the post-selection library for binding to *parS*.

N wt, *parS* post-selection library = counts of the WT (RTAG) in the post selection library for binding to *parS*.

N variant, pre-selection library = counts of each variant in the pre-selection (starting) library.

N wt, pre-selection library = counts of the WT (RTAG) in the pre-selection (starting) library.

**f**_*NBS, raw*_ *= log*_*10*_*(N variant, NBS post-selection library / N wt, NBS post-selection library) - log*_*10*_*(N variant, pre-selection library / N wt, pre-selection library)*

N variant, *NBS* post-selection library = counts of each variant in the post-selection library for binding to *NBS*.

N wt, *NBS* post-selection library = counts of the WT (QKKR) in the post selection library for binding to *NBS*.

N variant, pre-selection library = counts of each variant in the pre-selection (starting) library.

N wt, pre-selection library = counts of the WT (QKKR) in the pre-selection (starting) library.

These raw fitness scores were further transformed so that **f**_*parS*_ of the RTAG variant was 1 and that of QKKR variant was 0, and **f**_*NBS*_ of the RTAG variant was 0 and that of QKKR variant was 1. The fitness scores for every variant were presented in the fitness scatterplot (Figure 5C). Dark green: strong *parS* binding, no *NBS* binding (fitness score: *f*_*parS*_≥0.6, *f*_*NBS*_≤0.2); light green: strong *parS* binding, weak-to-medium *NBS* binding (*f*_*parS*_≥0.6, 0.2≤*f*_*NBS*_≤0.6); magenta: strong *NBS* binding, no *parS* binding (*f*_*NBS*_≥0.6, *f*_*parS*_≤0.2); pink: strong *NBS* binding, weak-to-medium *parS* binding (*f*_*NBS*_≥0.6, 0.2≤*f*_*parS*_≤0.6); black: dual specificity i.e. bind strongly to both *parS* and *NBS* (*f*_*NBS*_≥0.6 *f*_*parS*_≥0.6). Frequency logos of each class of variants were constructed using WebLogo 3.0 [42]

#### Reproducibility among replicates

To check the reproducibility among replicates, we plotted log_10_(counts) of each variant in replicate 1 vs. replicate 2 (and vs. replicate 3). Only variants with more than four reads were included in such plot. We used R to calculate Pearson’s correlation coefficients (*R*^*2*^) and to plot the linear best fit (Figure S5C). We found that independent experiments were reproducible (*R*^*2*^=0.86-0.98) (Figure S5C). Reads from three independent replicates were subsequently pooled together for the pre-selection, *parS* post-selection, and *NBS* post-selection experiments. Pooled reads were used to construct the scatterplot and frequency sequence logos (Figure 5C), and for the construction of the network graph (Figure 6B).

#### Generation of force-directed networks graphs and analysis of shortest paths

We constructed a force-directed graph that connects functional variants (nodes) together by lines (edges) if they are different by a single amino acid (AA) (Figure 6B). The node size is proportional to its connectivity (number of edges), and node colors represent different classes of functional variants (Figure 6B). Similarly, we also created a network graph in which edges represent variants that differ by a single nucleotide (nt) substitution, following a standard codon table (Figure S7A). Force-directed graphs were generated using Gephi network visualization software. Node and edge files were prepared in R. The network layout was generated by running the ForceAtlas algorithm that was implemented in Gephi. Default parameters were used for the ForceAtlas algorithm, except that the repulsion and attraction strength were set to 200 and 10, respectively. The ForceAtlas algorithm arranged nodes in the two-dimensional space based on connectivity: nodes tend to repel each other but they are attracted to each other if these exists a connectivity (an edge). As the result of running the Force Atlas to completion, densely interconnected nodes are clustered together while less well-connected nodes are forced to different spatial locations. To analyze the properties of the network and the mutational paths that traverse the network, we employed the *igraph* package implemented in R. Our network did not include non-functional (grey) variants/nodes.

## QUANTIFICATION AND STATISTICAL ANALYSIS

Information about statistical analysis and sample size for each experiment are detailed in the relevant Methods sections.

---

**Figure S1.**
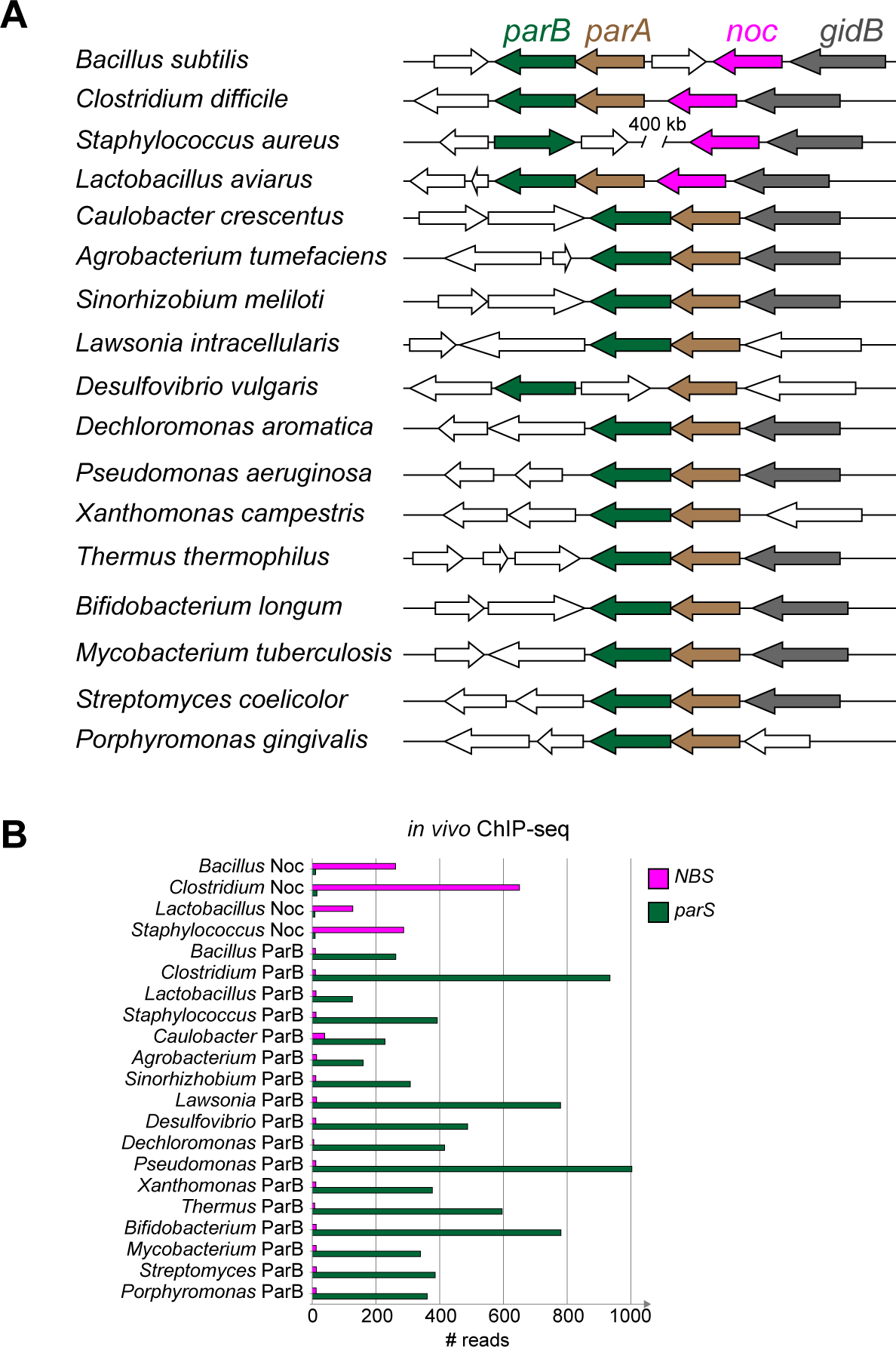
DNA-binding specificity for *parS* and *NBS* is conserved among ParB and Noc orthologs, related to Figure 1. **(A)** Genomic context of ParB- and Noc-encoding genes in various bacterial species. *parB, parA, noc*, and the highly conserved *gidB* gene, are colored in dark green, brown, magenta, and grey, respectively. Genes at the border of the *parB-parA-noc* cluster (open arrows) vary between bacterial species. **(B)** The *in vivo* binding preferences of ParB/Noc to *parS/NBS* as measured by ChIP-seq. An *E. coli* strain with a single *parS* and *NBS* site engineered onto the chromosome was used as a heterologous host for expression of FLAG-tagged ParB/Noc. For ChIP-seq data, reads in a 100-bp window surrounding the *parS/NBS* site were quantified and used as a proxy for the enrichment of immunoprecipitated *parS* or *NBS* DNA.

**Figure S2.**
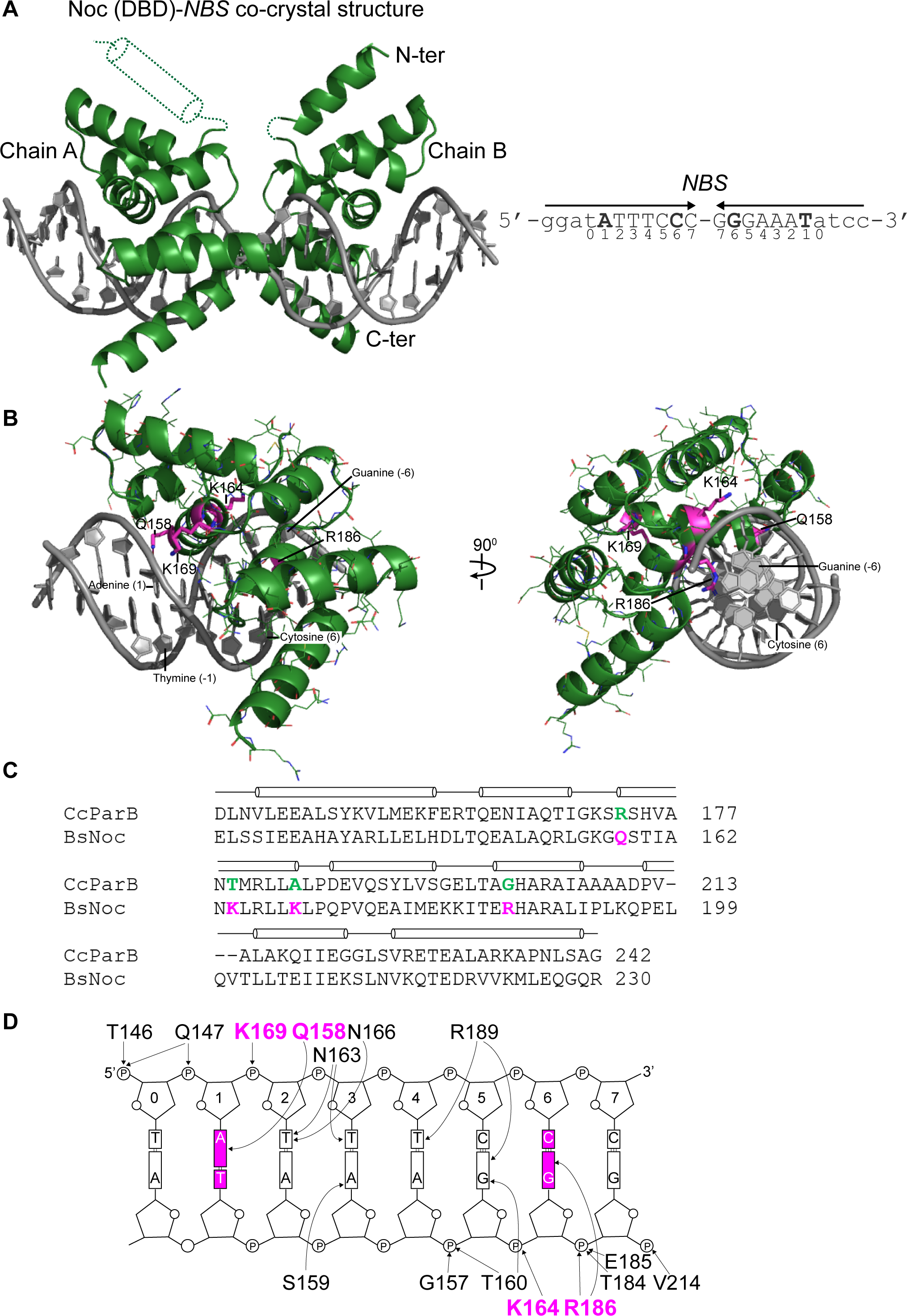
Co-crystal structures of the Noc (DBD)-*NBS* complex, related to Figure 4. **(A)** The structure of two Noc (DNA-binding domain) monomers (dark green) in complex with a 22-bp *NBS* DNA (grey). A helix (residues 113-125, dotted dark green cylinder) in chain A is not resolved. The nucleotide sequence of the 22-bp *NBS* site is shown on the left-hand side; bases (Adenine 1 and Cytosine 6) that are different from *parS* are in bold. **(B)** One monomer of Noc (DBD) is shown in complex with an *NBS* half-site; four core specificity residues are shown in stick presentation, labeled, and colored in magenta. Other residues surrounding specificity residues are showed as lines and colored in dark green. **(C)** Amino acid sequences of *C. crescentus* ParB and *B. subtilis* Noc with the positions of four specificity residues highlighted in dark green and magenta, respectively. Secondary structures are shown above the sequence alignment. **(D)** Schematic representation of Noc (DBD)-*NBS* interactions. For simplicity, only half of *NBS* is shown. The two bases at position 1 and 6 that are different between *parS* and *NBS* are highlighted in magenta. The four core specificity residues are also colored in magenta.

**Figure S3.**
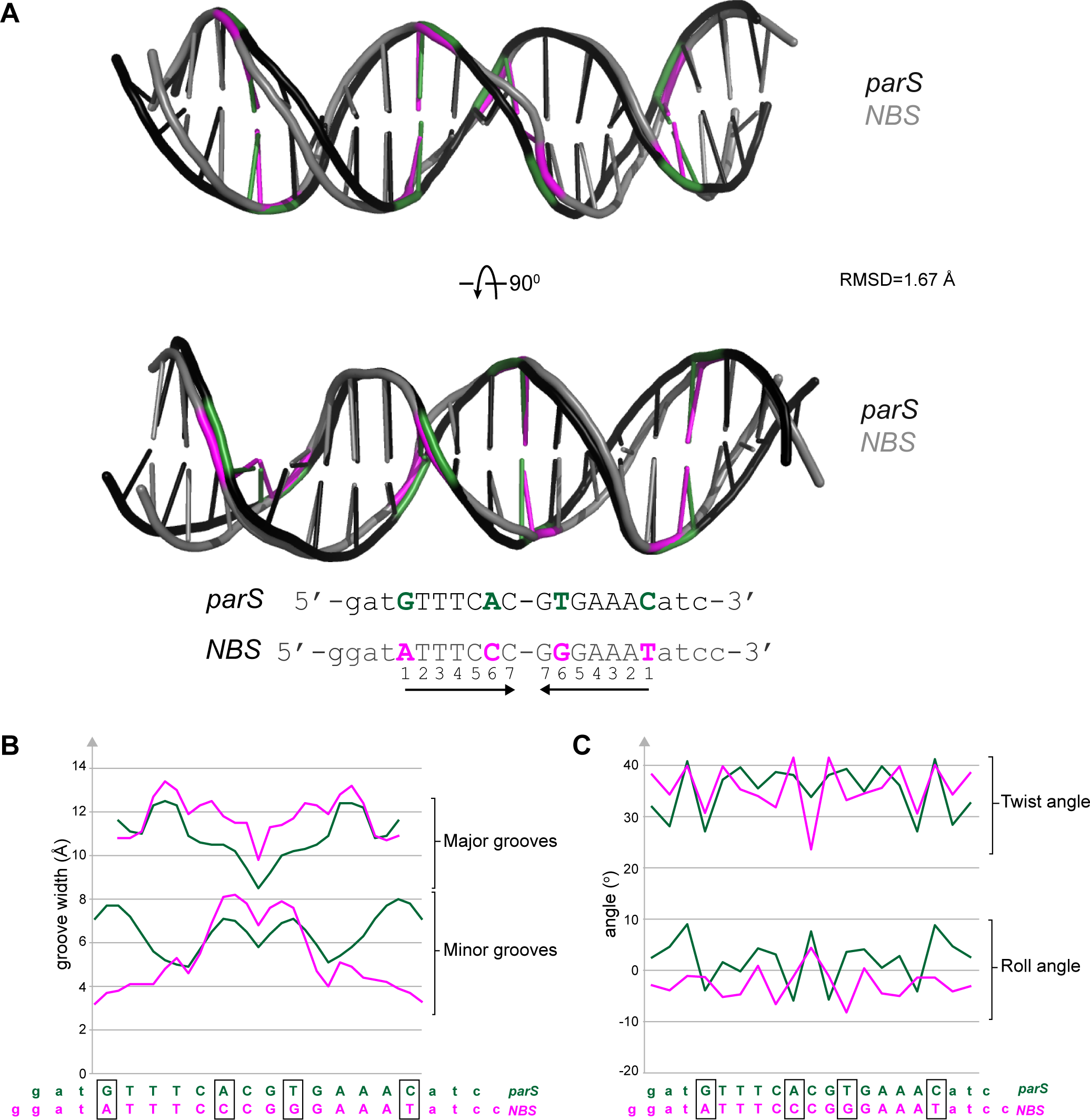
Conformational changes at *parS* and *NBS* DNA within the two co-crystal structures, related to Figure 4. **(A)** Superimposition of *parS* and *NBS* DNA structures, root-mean-square deviation (RMSD) value is also shown. Bases that differ between *parS* (dark green) and *NBS* (magenta) are highlighted. **(B)** The major and minor groove widths of the bound DNA (*parS*: dark green, *NBS*: magenta). **(C)** The roll and twist angles for each base pair step of the bound DNA (*parS*: dark green, *NBS*: magenta).

**Figure S4.**
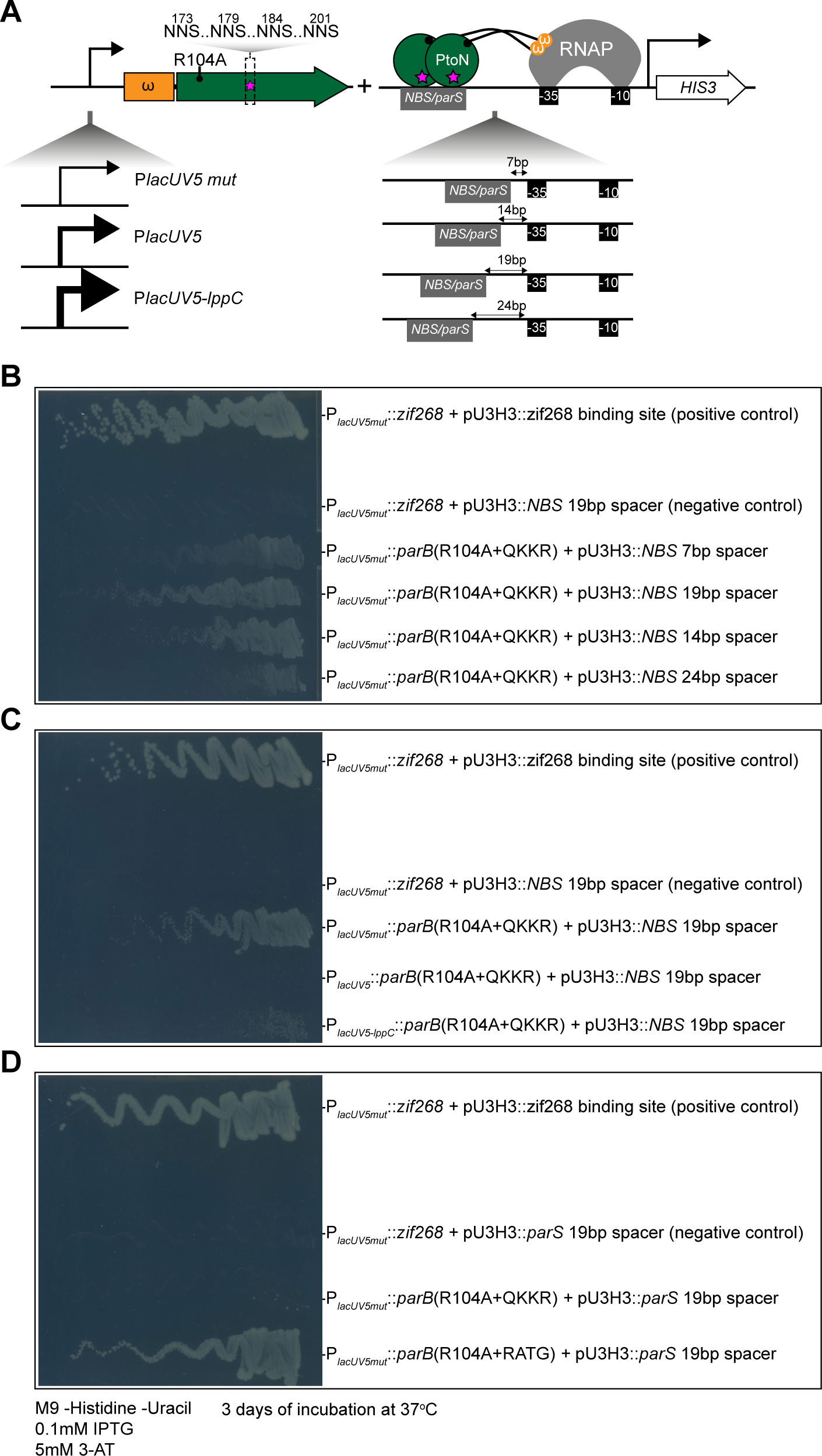
Optimization of bacterial one-hybrid (B1H) assay to select for variants that bind *parS* or *NBS*, related to Figure 5. **(A)** The strength of the promoter that drives the expression of *parB* variants and the distances between the *parS* or *NBS* binding site to the core −10 −35 promoter were optimized. **(B)** A 19-bp gap between *NBS/parS* and the core promoter is optimal, based on the streak test for cell growth in a minimal medium lacking histidine. **(C)** A weak promoter (P_*lacUV5mut*_) is optimal, based on the streak test for cell growth in a minimal medium lacking histidine. **(B and D)** The presence of *parS* or *NBS* upstream of HIS3, in the absence of ParB variants, did not auto-activate its expression.

**Figure S5.**
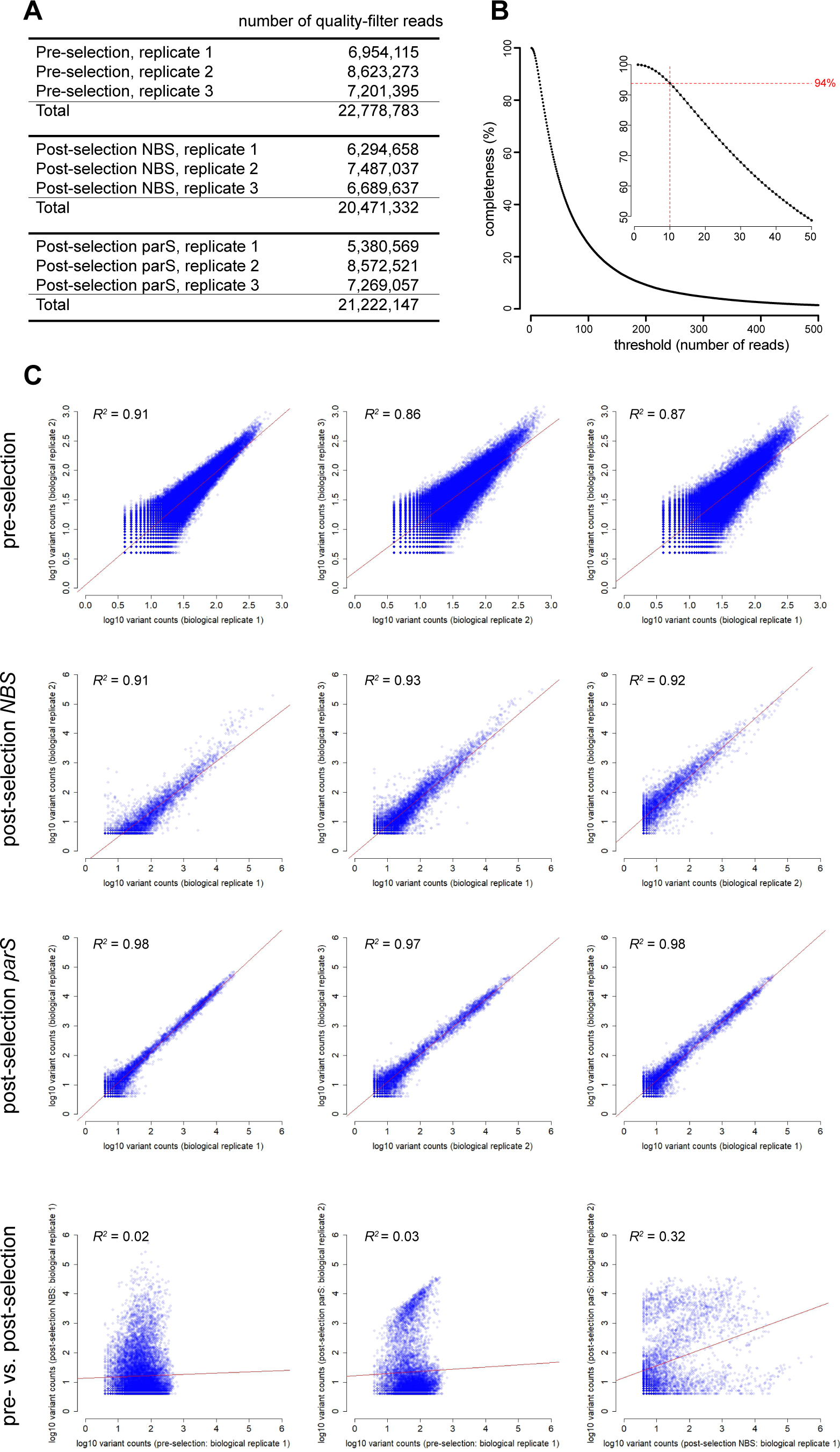
Statistics of deep sequencing reads and completeness of starting libraries, related to Figure 5. **(A)** Number of quality-filtered reads for each biological replicate, for pre- and post-selection libraries. **(B)** The completeness of pre-selection libraries at different thresholds. A completeness of 100% means that all 160,000 variants lacking stop codons were present in the pre-selection library. In the starting library, greater than 94% of the predicted variants were represented by at least 10 reads. **(C)** Reproducibility of biological replicates: pre- vs. pre-selection replicates and pre- vs. post-selection replicates. Pearson’s correlation coefficients (*R*^*2*^) are also shown. Red lines show least squares best fits.

**Figure S6.**
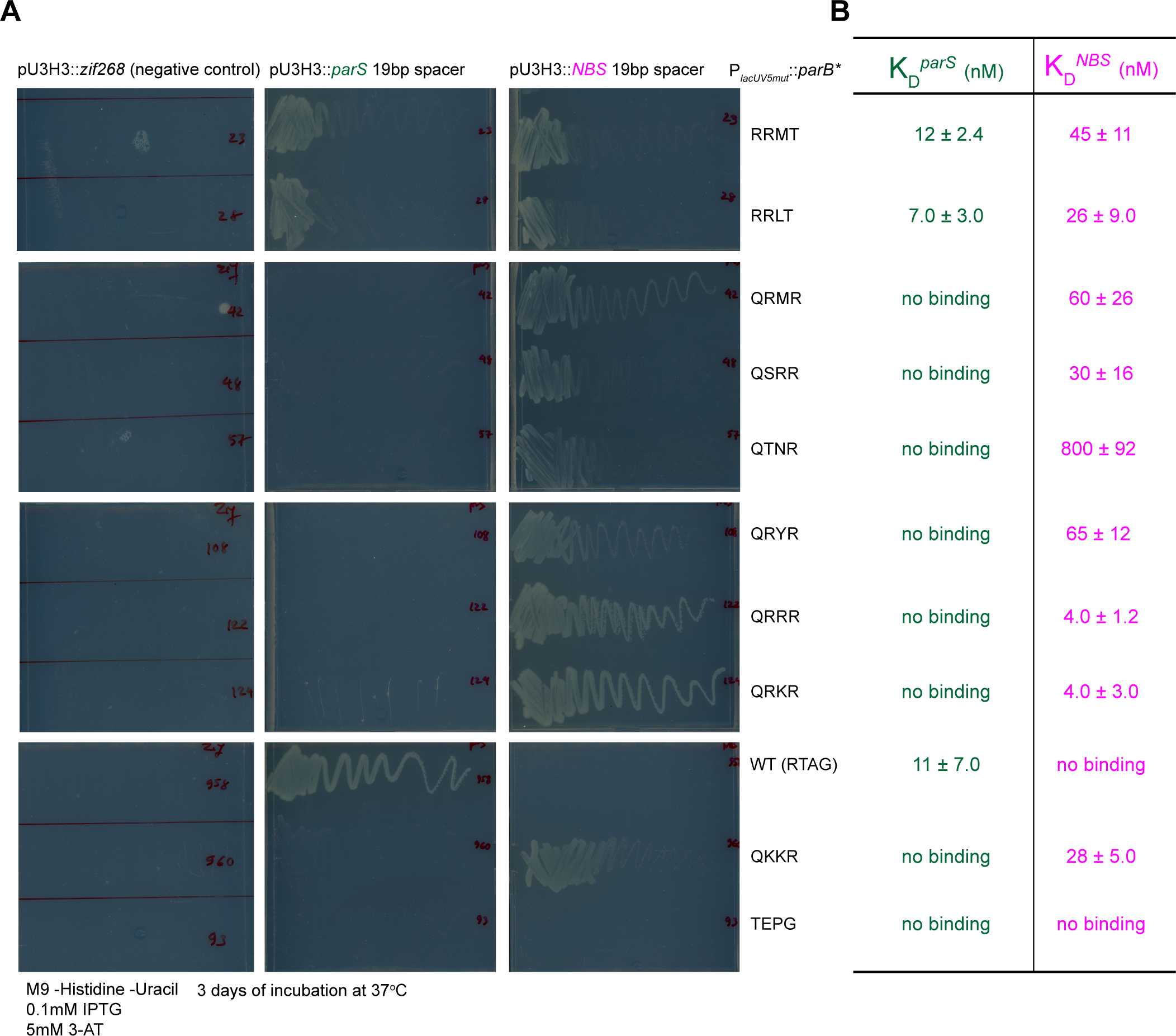
Validation of selected variants from the deep mutational scanning experiments, related to Figure 5. **(A)** Validation by pairwise bacterial one-hybrid assays. The ability of nine selected variants to grow on a minimal medium lacking histidine (but supplemented with 5mM 3-AT to increase the stringency) was assessed by a streak test. Plasmid harboring a binding site of an eukaryotic transcription factor (*zif268*) served as a negative control. **(B)** Validation by bio-layer interferometry assays. Selected variants were expressed, purified, and K_D_ ± SD were measured by bio-layer interferometry assay.

**Figure S7.**
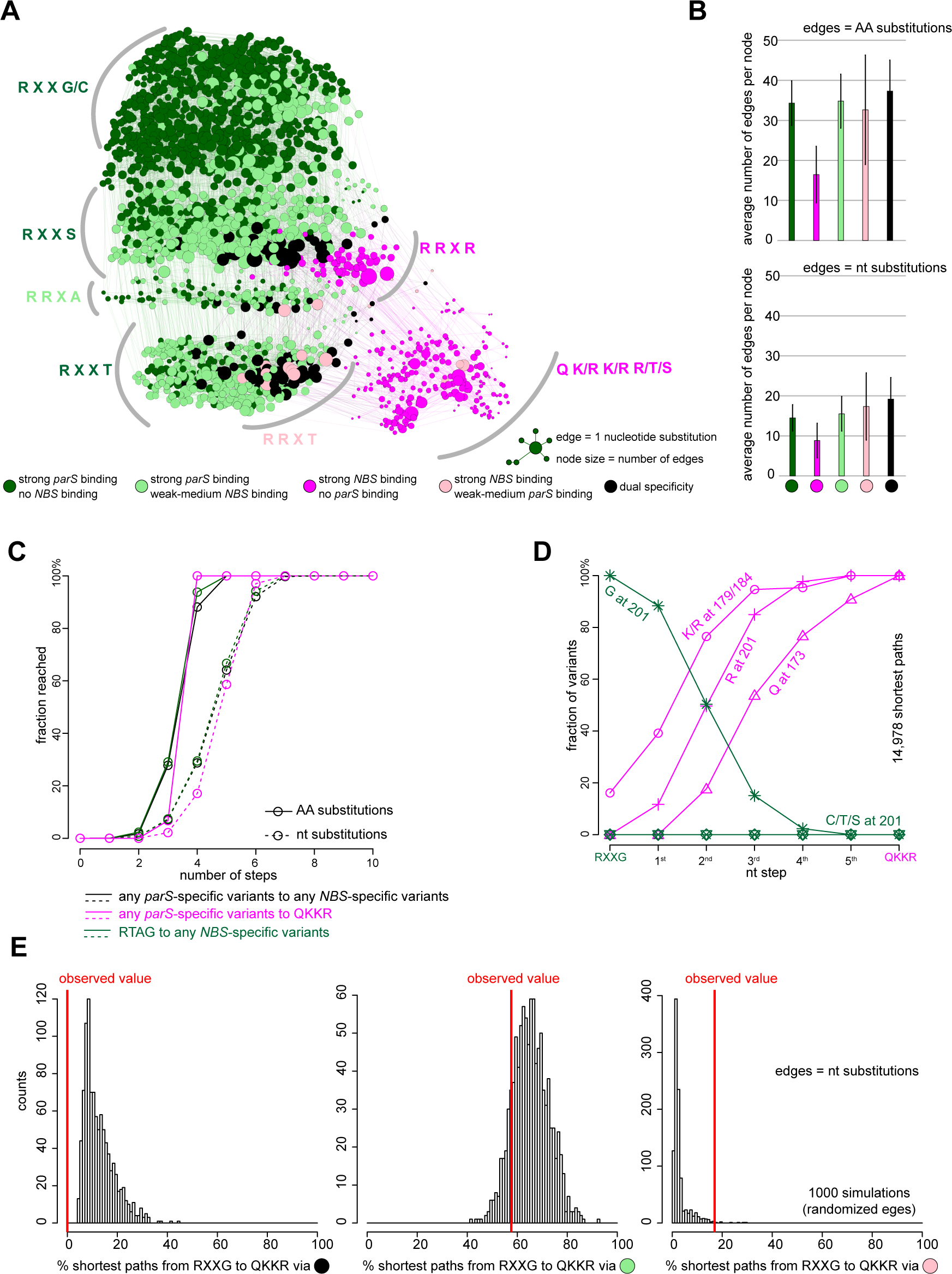
Deep mutational scanning experiments reveal the common properties of mutational paths, related to Figure 6. **(A)** A force-directed network graph connecting strong *parS*-binding variants to strong *NBS*-binding variants. Nodes represent individual variants, and edges represent single nucleotide (nt) substitutions. Node sizes are proportional to their corresponding numbers of edges. Node colors correspond to different classes of variants. **(B)** Average number of edges per node. **(C)** Cumulative fraction of variants that reached their destinations in a given number of amino acid (solid line) or nucleotide (dotted line) substitutions. Black lines: from any *parS*-specific variants to any *NBS*-specific variants. Magenta lines: from any *parS*-specific variants to QKKR. Dark green lines: from RTAG to any *NBS*-specific variants. **(D)** Fraction of intermediates on all shortest paths from highly *parS*-specific RXXG variants to the *NBS*-preferred QKKR that have permissive amino acids (K/R) at either position 179/184 or both, or have R at position 201, or Q at position 173, or C/T/S at position 201 after a given number of nt steps. **(E)** Percentage of shortest paths that traversed black, light green, or pink variants to reach QKKR from any of the highly *parS*-specific RXXG variants (red lines). The result was compared to ones from 1,000 simulations where the by-nt-substitution edges were shuffled randomly while keeping the total number of nodes, edges, and graph density constant.

## Notes

### Competing Interest Statement

The authors have declared no competing interest.

### Summary of Updates

1) We have now measured KD for proteins used in this study (Fig. 2D, Fig. 3A and C-F, and Fig. S6B), and have updated the Results, Figures, and Materials&Methods sections accordingly. 2) We were able to crystallize and subsequently solve the structure of a wild-type Bacillus Noc (DNA-binding domain) in complex with a 22-bp NBS DNA duplex (Fig. S2 and Fig. 4, Table S4 for data collection and processing statistics, Materials&Methods have been updated).

## REFERENCES

1. Kaessmann H. 2010 Origins, evolution, and phenotypic impact of new genes. Genome Res. 20, 1313–1326. (doi: 10.1101/gr.101386.109)

2. Qian W, Zhang J. 2014 Genomic evidence for adaptation by gene duplication. Genome Res 24, 1356–1362. (doi: 10.1101/gr.172098.114)

3. Conrad B, Antonarakis SE. 2007 Gene duplication: a drive for phenotypic diversity and cause of human disease. Annu Rev Genomics Hum Genet 8, 17–35. (doi: 10.1146/annurev.genom.8.021307.110233)

4. Lynch M, Conery JS. 2000 The evolutionary fate and consequences of duplicate genes. Science 290, 1151–1155.

5. Teichmann SA, Babu MM. 2004 Gene regulatory network growth by duplication. Nat. Genet. 36, 492–496. (doi: 10.1038/ng1340)

6. Livny J, Yamaichi Y, Waldor MK. 2007 Distribution of Centromere-Like parS Sites in Bacteria: Insights from Comparative Genomics. J. Bacteriol. 189, 8693–8703. (doi: 10.1128/JB.01239-07)

7. Lin DC-H, Grossman AD. 1998 Identification and Characterization of a Bacterial Chromosome Partitioning Site. Cell 92, 675–685. (doi: 10.1016/S0092-8674(00)81135-6)

8. Lagage V, Boccard F, Vallet-Gely I. 2016 Regional Control of Chromosome Segregation in Pseudomonas aeruginosa. PLOS Genetics 12, e1006428. (doi: 10.1371/journal.pgen.1006428)

9. Lim HC, Surovtsev IV, Beltran BG, Huang F, Bewersdorf J, Jacobs-Wagner C. 2014 Evidence for a DNA-relay mechanism in ParABS-mediated chromosome segregation. Elife 3, e02758. (doi: 10.7554/eLife.02758)

10. Toro E, Hong S-H, McAdams HH, Shapiro L. 2008 Caulobacter requires a dedicated mechanism to initiate chromosome segregation. PNAS 105, 15435–15440. (doi: 10.1073/pnas.0807448105)

11. Fisher GL et al. 2017 The structural basis for dynamic DNA binding and bridging interactions which condense the bacterial centromere. Elife 6. (doi: 10.7554/eLife.28086)

12. Fogel MA, Waldor MK. 2006 A dynamic, mitotic-like mechanism for bacterial chromosome segregation. Genes Dev. 20, 3269–3282. (doi: 10.1101/gad.1496506)

13. Gruber S, Errington J. 2009 Recruitment of condensin to replication origin regions by ParB/SpoOJ promotes chromosome segregation in B. subtilis. Cell 137, 685–696. (doi: 10.1016/j.cell.2009.02.035)

14. Ireton K, Gunther NW, Grossman AD. 1994 spo0J is required for normal chromosome segregation as well as the initiation of sporulation in Bacillus subtilis. J. Bacteriol. 176, 5320–5329. (doi: 10.1128/jb.176.17.5320-5329.1994)

15. Mohl DA, Gober JW. 1997 Cell cycle-dependent polar localization of chromosome partitioning proteins in Caulobacter crescentus. Cell 88, 675–684. (doi: 10.1016/s0092-8674(00)81910-8)

16. Tran NT, Laub MT, Le TBK. 2017 SMC Progressively Aligns Chromosomal Arms in Caulobacter crescentus but Is Antagonized by Convergent Transcription. Cell Rep 20, 2057–2071. (doi: 10.1016/j.celrep.2017.08.026)

17. Tran NT, Stevenson CE, Som NF, Thanapipatsiri A, Jalal ASB, Le TBK. 2018 Permissive zones for the centromere-binding protein ParB on the Caulobacter crescentus chromosome. Nucleic Acids Res 46, 1196–1209. (doi: 10.1093/nar/gkx1192)

18. Wang X, Le TBK, Lajoie BR, Dekker J, Laub MT, Rudner DZ. 2015 Condensin promotes the juxtaposition of DNA flanking its loading site in Bacillus subtilis. Genes Dev. 29, 1661–1675. (doi: 10.1101/gad.265876.115)

19. Harms A, Treuner-Lange A, Schumacher D, Sogaard-Andersen L. 2013 Tracking of chromosome and replisome dynamics in Myxococcus xanthus reveals a novel chromosome arrangement. PLoS Genet 9, e1003802. (doi: 10.1371/journal.pgen.1003802)

20. Jakimowicz D, Chater K, Zakrzewska-Czerwínska J. 2002 The ParB protein of Streptomyces coelicolor A3(2) recognizes a cluster of parS sequences within the origin-proximal region of the linear chromosome. Molecular Microbiology 45, 1365–1377. (doi: 10.1046/j.1365-2958.2002.03102.x)

21. Kawalek A, Bartosik AA, Glabski K, Jagura-Burdzy G. 2018 Pseudomonas aeruginosa partitioning protein ParB acts as a nucleoid-associated protein binding to multiple copies of a parS-related motif. Nucleic Acids Res. 46, 4592–4606. (doi: 10.1093/nar/gky257)

22. Murray H, Ferreira H, Errington J. 2006 The bacterial chromosome segregation protein Spo0J spreads along DNA from parS nucleation sites. Molecular Microbiology 61, 1352–1361. (doi: 10.1111/j.1365-2958.2006.05316.x)

23. Sievers J, Raether B, Perego M, Errington J. 2002 Characterization of the parB-Like yyaA Gene of Bacillus subtilis. J. Bacteriol. 184, 1102–1111. (doi: 10.1128/jb.184.4.1102-1111.2002)

24. Wu LJ, Errington J. 2004 Coordination of cell division and chromosome segregation by a nucleoid occlusion protein in Bacillus subtilis. Cell 117, 915–925. (doi: 10.1016/j.cell.2004.06.002)

25. Wu LJ, Ishikawa S, Kawai Y, Oshima T, Ogasawara N, Errington J. 2009 Noc protein binds to specific DNA sequences to coordinate cell division with chromosome segregation. EMBO J. 28, 1940–1952. (doi: 10.1038/emboj.2009.144)

26. Pang T, Wang X, Lim HC, Bernhardt TG, Rudner DZ. 2017 The nucleoid occlusion factor Noc controls DNA replication initiation in Staphylococcus aureus. PLOS Genetics 13, e1006908. (doi: 10.1371/journal.pgen.1006908)

27. Wu LJ, Errington J. 2011 Nucleoid occlusion and bacterial cell division. Nat. Rev. Microbiol. 10, 8–12. (doi: 10.1038/nrmicro2671)

28. Noyes MB, Meng X, Wakabayashi A, Sinha S, Brodsky MH, Wolfe SA. 2008 A systematic characterization of factors that regulate Drosophila segmentation via a bacterial one-hybrid system. Nucleic Acids Res 36, 2547–2560. (doi: 10.1093/nar/gkn048)

29. Lee PS, Grossman AD. 2006 The chromosome partitioning proteins Soj (ParA) and Spo0J (ParB) contribute to accurate chromosome partitioning, separation of replicated sister origins, and regulation of replication initiation in Bacillus subtilis. Mol. Microbiol. 60, 853–869. (doi: 10.1111/j.1365-2958.2006.05140.x)

30. Podgornaia AI, Laub MT. 2015 Protein evolution. Pervasive degeneracy and epistasis in a protein-protein interface. Science 347, 673–677. (doi: 10.1126/science.1257360)

31. Bloom JD, Gong LI, Baltimore D. 2010 Permissive Secondary Mutations Enable the Evolution of Influenza Oseltamivir Resistance. Science 328, 1272–1275. (doi: 10.1126/science.1187816)

32. Gong LI, Suchard MA, Bloom JD. 2013 Stability-mediated epistasis constrains the evolution of an influenza protein. eLife 2, e00631. (doi: 10.7554/eLife.00631)

33. Wang X, Minasov G, Shoichet BK. 2002 Evolution of an antibiotic resistance enzyme constrained by stability and activity trade-offs. J. Mol. Biol. 320, 85–95. (doi: 10.1016/S0022-2836(02)00400-X)

34. McKeown AN, Bridgham JT, Anderson DW, Murphy MN, Ortlund EA, Thornton JW. 2014 Evolution of DNA specificity in a transcription factor family produced a new gene regulatory module. Cell 159, 58–68. (doi: 10.1016/j.cell.2014.09.003)

35. Starr TN, Picton LK, Thornton JW. 2017 Alternative evolutionary histories in the sequence space of an ancient protein. Nature 549, 409. (doi: 10.1038/nature23902)

36. Aakre CD, Herrou J, Phung TN, Perchuk BS, Crosson S, Laub MT. 2015 Evolving New Protein-Protein Interaction Specificity through Promiscuous Intermediates. Cell 163, 594–606. (doi: 10.1016/j.cell.2015.09.055)

37. Raumann BE, Knight KL, Sauer RT. 1995 Dramatic changes in DNA-binding specificity caused by single residue substitutions in an Arc/Mnt hybrid repressor. Nature Structural & Molecular Biology 2, 1115–1122. (doi: 10.1038/nsb1295-1115)

38. Ivankov DN, Finkelstein AV, Kondrashov FA. 2014 A structural perspective of compensatory evolution. Current Opinion in Structural Biology 26, 104–112. (doi: 10.1016/j.sbi.2014.05.004)

39. Sikosek T, Chan HS. 2014 Biophysics of protein evolution and evolutionary protein biophysics. J R Soc Interface 11, 20140419. (doi: 10.1098/rsif.2014.0419)

40. Starr TN, Thornton JW. 2016 Epistasis in protein evolution. Protein Sci 25, 1204–1218. (doi: 10.1002/pro.2897)

41. Steinegger M, Meier M, Mirdita M, Voehringer H, Haunsberger SJ, Soeding J. 2019 HH-suite3 for fast remote homology detection and deep protein annotation. bioRxiv, 560029. (doi: 10.1101/560029)

42. Crooks GE, Hon G, Chandonia J-M, Brenner SE. 2004 WebLogo: a sequence logo generator. Genome Res. 14, 1188–1190. (doi: 10.1101/gr.849004)

43. Letunic I, Bork P. 2016 Interactive tree of life (iTOL) v3: an online tool for the display and annotation of phylogenetic and other trees. Nucleic Acids Res. 44, W242–245. (doi: 10.1093/nar/gkw290)

44. Miller MA, Pfeiffer W, Schwartz T. 2011 The CIPRES Science Gateway: A Community Resource for Phylogenetic Analyses. In Proceedings of the 2011 TeraGrid Conference: Extreme Digital Discovery, pp. 41:1–41:8. New York, NY, USA: ACM. (doi: 10.1145/2016741.2016785)

45. Lavery R, Moakher M, Maddocks JH, Petkeviciute D, Zakrzewska K. 2009 Conformational analysis of nucleic acids revisited: Curves+. Nucleic Acids Res 37, 5917–5929. (doi: 10.1093/nar/gkp608)

46. Langmead B, Trapnell C, Pop M, Salzberg SL. 2009 Ultrafast and memory-efficient alignment of short DNA sequences to the human genome. Genome Biol 10, R25.

47. Quinlan AR, Hall IM. 2010 BEDTools: a flexible suite of utilities for comparing genomic features. Bioinformatics 26, 841–842. (doi: 10.1093/bioinformatics/btq033)

48. Winter G et al. 2018 DIALS: implementation and evaluation of a new integration package. Acta Crystallogr D Struct Biol 74, 85–97. (doi: 10.1107/S2059798317017235)

49. Evans PR, Murshudov GN. 2013 How good are my data and what is the resolution? Acta Cryst D 69, 1204–1214. (doi: 10.1107/S0907444913000061)

50. Potterton L et al. 2018 CCP4i2: the new graphical user interface to the CCP4 program suite. Acta Crystallogr D Struct Biol 74, 68–84. (doi: 10.1107/S2059798317016035)

51. Emsley P, Cowtan K. 2004 Coot: model-building tools for molecular graphics. Acta Crystallogr. D Biol. Crystallogr. 60, 2126–2132. (doi: 10.1107/S0907444904019158)

52. McCoy AJ, Grosse-Kunstleve RW, Adams PD, Winn MD, Storoni LC, Read RJ. 2007 Phaser crystallographic software. J Appl Crystallogr 40, 658–674. (doi: 10.1107/S0021889807021206)

53. Murshudov GN, Vagin AA, Dodson EJ. 1997 Refinement of Macromolecular Structures by the Maximum-Likelihood Method. Acta Cryst D 53, 240–255. (doi: 10.1107/S0907444996012255)

54. Cowtan K. 2006 The Buccaneer software for automated model building. 1. Tracing protein chains. Acta Crystallogr. D Biol. Crystallogr. 62, 1002–1011. (doi: 10.1107/S0907444906022116)

55. Chen VB, Arendall WB, Headd JJ, Keedy DA, Immormino RM, Kapral GJ, Murray LW, Richardson JS, Richardson DC. 2010 MolProbity: all-atom structure validation for macromolecular crystallography. Acta Crystallogr. D Biol. Crystallogr. 66, 12–21. (doi: 10.1107/S0907444909042073)

56. Krissinel E. 2015 Stock-based detection of protein oligomeric states in jsPISA. Nucleic Acids Res. 43, W314–319. (doi: 10.1093/nar/gkv314)

57. Karimova G, Pidoux J, Ullmann A, Ladant D. 1998 A bacterial two-hybrid system based on a reconstituted signal transduction pathway. Proc Natl Acad Sci U S A 95, 5752–5756.

58. Edgar RC. 2004 MUSCLE: multiple sequence alignment with high accuracy and high throughput. Nucleic Acids Res. 32, 1792–1797. (doi: 10.1093/nar/gkh340)

59. Stamatakis A. 2014 RAxML version 8: a tool for phylogenetic analysis and post–analysis of large phylogenies. Bioinformatics 30, 1312–1313. (doi: 10.1093/bioinformatics/btu033)

60. Winter G. 2010 xia2: an expert system for macromolecular crystallography data reduction. J Appl Cryst 43, 186–190. (doi: 10.1107/S0021889809045701)

61. Chen B-W, Lin M-H, Chu C-H, Hsu C-E, Sun Y-J. 2015 Insights into ParB spreading from the complex structure of Spo0J and parS. Proc. Natl. Acad. Sci. U.S.A. 112, 6613–6618. (doi: 10.1073/pnas.1421927112)

62. Bunkóczi G, Read RJ. 2011 Improvement of molecular-replacement models with Sculptor. Acta Crystallogr D Biol Crystallogr 67, 303–312. (doi: 10.1107/S0907444910051218)

63. Anandakrishnan R, Aguilar B, Onufriev AV. 2012 H++ 3.0: automating pK prediction and the preparation of biomolecular structures for atomistic molecular modeling and simulations. Nucleic Acids Res. 40, W537–541. (doi: 10.1093/nar/gks375)

64. Maier JA, Martinez C, Kasavajhala K, Wickstrom L, Hauser KE, Simmerling C. 2015 ff14SB: Improving the Accuracy of Protein Side Chain and Backbone Parameters from ff99SB. J Chem Theory Comput 11, 3696–3713. (doi: 10.1021/acs.jctc.5b00255)

65. Ivani I et al. 2016 Parmbsc1: a refined force field for DNA simulations. Nat. Methods 13, 55–58. (doi: 10.1038/nmeth.3658)

66. Price DJ, Brooks CL. 2004 A modified TIP3P water potential for simulation with Ewald summation. J Chem Phys 121, 10096–10103. (doi: 10.1063/1.1808117)

67. Smith DE, Dang LX. 1994 Computer simulations of NaCl association in polarizable water. J. Chem. Phys. 100, 3757–3766. (doi: 10.1063/1.466363)

68. Noy A, Golestanian R. 2010 The chirality of DNA: elasticity cross-terms at base-pair level including A-tracts and the influence of ionic strength. J Phys Chem B 114, 8022–8031. (doi: 10.1021/jp104133j)

69. Ryckaert J-P, Ciccotti G, Berendsen HJC. 1977 Numerical integration of the cartesian equations of motion of a system with constraints: molecular dynamics of n-alkanes. Journal of Computational Physics 23, 327–341. (doi: 10.1016/0021-9991(77)90098-5)

70. Darden T, York D, Pedersen L. 1993 Particle mesh Ewald: An N·log(N) method for Ewald sums in large systems. J. Chem. Phys. 98, 10089–10092. (doi: 10.1063/1.464397)

71. Roe DR, Cheatham TE. 2013 PTRAJ and CPPTRAJ: Software for Processing and Analysis of Molecular Dynamics Trajectory Data. J. Chem. Theory Comput. 9, 3084–3095. (doi: 10.1021/ct400341p)

72. Chen C, Esadze A, Zandarashvili L, Nguyen D, Pettitt BM, Iwahara J. 2015 Dynamic Equilibria of Short-Range Electrostatic Interactions at Molecular Interfaces of Protein–DNA Complexes. J Phys Chem Lett 6, 2733–2737. (doi: 10.1021/acs.jpclett.5b01134)

73. Bosley AD, Ostermeier M. 2005 Mathematical expressions useful in the construction, description and evaluation of protein libraries. Biomol. Eng. 22, 57–61. (doi: 10.1016/j.bioeng.2004.11.002)

74. van Opijnen T, Bodi KL, Camilli A. 2009 Tn-seq: high-throughput parallel sequencing for fitness and genetic interaction studies in microorganisms. Nat. Methods 6, 767–772. (doi: 10.1038/nmeth.1377)

