## Supplementary for "Diversification of DNA-binding specificity via permissive and specificity-switching mutations in the ParB/Noc protein family"

### TABLE S1. STRAINS

| **Strains** | **Strains/descriptions** | **Source** |
|---|---|---|
| AB1157 | *thr-1, ara-14, leuB6, Δ(gpt-proA)62, lacY1, tsx-33, supE44, galK2, rac-, hisG4(Oc), rfbD1, mgl-51, rpsL31, kdgK51, xyl-5, mtl-1, argE3 (Oc), thi-1, qsr-* | Yale *E. coli* Genetic Stock Center |
| DH5α | *E. coli* host for DNA cloning and propagation of plasmid | Le lab collection |
| Rosetta (DE3) | *E. coli* host for protein overexpression from an IPTG-inducible T7 promoter | Merck |
| CJW4025 | BL21 pET21b::*parB-(his)6* | Gift from Christine Jacob-Wagner (Lim et al., 2014) |
| TLE3000 | *AB1157 ybbD::parS::markerless ygcE::NBS::markerless* | This study |
| TLE3001 | *USO rpoZ- hisB- pyrF-* | Scott Wolfe (Noyes et al., 2008) via Addgene |
| | TLE3000+pUT18C::1xFLAG-*Bacillus subtilis* Noc | This study |
| | TLE3000+pUT18C::1xFLAG-*Clostridium difficile* Noc | This study |
| | TLE3000+pUT18C::1xFLAG-*Lactobacillus aviarius* Noc | This study |
| | TLE3000+pUT18C::1xFLAG-*Staphylococcus aureus* Noc | This study |
| | TLE3000+pUT18C::1xFLAG-*Bacillus subtilis* ParB | This study |
| | TLE3000+pUT18C::1xFLAG-*Clostridium difficile* ParB | This study |
| | TLE3000+pUT18C::1xFLAG-*Lactobacillus aviarius* ParB | This study |
| | TLE3000+pUT18C::1xFLAG-*Staphylococcus aureus* ParB | This study |
| | TLE3000+pUT18C::1xFLAG-*Caulobacter crescentus* ParB | This study |
| | TLE3000+pUT18C::1xFLAG-*Agrobacterium tumefaciens* ParB | This study |
| | TLE3000+pUT18C::1xFLAG-*Sinorhizobium meliloti* ParB | This study |
| | TLE3000+pUT18C::1xFLAG-*Lawsonia intracellularis* ParB | This study |
| | TLE3000+pUT18C::1xFLAG-*Desulfovibrio vulgaris* ParB | This study |
| | TLE3000+pUT18C::1xFLAG-*Dechloromonas aromatica* ParB | This study |
| | TLE3000+pUT18C::1xFLAG-*Pseudomonas aeruginosa* ParB | This study |
| | TLE3000+pUT18C::1xFLAG-*Xanthomonas campestris* ParB | This study |
| | TLE3000+pUT18C::1xFLAG-*Thermus thermophilus* ParB | This study |
| | TLE3000+pUT18C::1xFLAG-*Bifidobacterium longum* ParB | This study |
| | TLE3000+pUT18C::1xFLAG-*Mycobacterium tuberculosis* ParB | This study |
| | TLE3000+pUT18C::1xFLAG-*Streptomyces coelicolor* ParB | This study |
| | TLE3000+pUT18C::1xFLAG-*Porphyromonas gingivalis* ParB | This study |
| | TLE3001 + pB1H2-w2::*zif268* + pU3H3::*zif268* binding site | This study |
| | TLE3001 + pB1H2-w2::*zif268* + pU3H3::19bp-*NBS* | This study |
| | TLE3001 + pB1H2-w2::*Caulobacter* ParB (R104A + Q173K179K184R201) + pU3H3::7bp-*NBS* | This study |
| | TLE3001 + pB1H2-w2::*Caulobacter* ParB (R104A + Q173K179K184R201) + pU3H3::14bp-*NBS* | This study |
| | TLE3001 + pB1H2-w2::*Caulobacter* ParB (R104A + Q173K179K184R201) + pU3H3::19bp-*NBS* | This study |
| | TLE3001 + pB1H2-w2::*Caulobacter* ParB (R104A + Q173K179K184R201) + pU3H3::24bp-*NBS* | This study |
| | TLE3001 + pB1H2-w5::*Caulobacter* ParB (R104A + | This study |

| | | |
|---|---|---|
| | Q173K179K184R201) + pU3H3::19bp-*NBS* | |
| | TLE3001 + pB1H2-w5L::*Caulobacter* ParB (R104A + Q173K179K184R201) + pU3H3::19bp-*NBS* | This study |
| | TLE3001 + pB1H2-w2::*zif268* + pU3H3::19bp-*parS* | This study |
| | TLE3001 + pB1H2-w2::*Caulobacter* ParB (R104A + Q173K179K184R201) + pU3H3::19bp-*parS* | This study |
| | TLE3001 + pB1H2-w2::*Caulobacter* ParB (R104A + R173A179T184G201) + pU3H3::19bp-*parS* | This study |
| | BL21 Rosetta pRARE + various pET21b-based protein overexpression vectors (see the plasmid list for the complete collection of protein overexpression plasmids) | This study |

### TABLE S2. PLASMIDS

| Plasmids | Description | Source |
|---|---|---|
| pENTR::D-TOPO | ENTRY vector for Gateway cloning, kanamycin$^R$ | Invitrogen |
| | pENTR::*Bacillus subtilis* Noc | This study |
| | pENTR::*Clostridium difficile* Noc | This study |
| | pENTR::*Lactobacillus aviarius* Noc | This study |
| | pENTR::*Staphylococcus aureus* Noc | This study |
| | pENTR::*Bacillus subtilis* ParB | This study |
| | pENTR::*Clostridium difficile* ParB | This study |
| | pENTR::*Lactobacillus aviarius* ParB | This study |
| | pENTR::*Staphylococcus aureus* ParB | This study |
| | pENTR::*Caulobacter crescentus* ParB | This study |
| | pENTR::*Agrobacterium tumefaciens* ParB | This study |
| | pENTR::*Sinorhizobium meliloti* ParB | This study |
| | pENTR::*Lawsonia intracellularis* ParB | This study |
| | pENTR::*Desulfovibrio vulgaris* ParB | This study |
| | pENTR::*Dechloromonas aromatica* ParB | This study |
| | pENTR::*Pseudomonas aeruginosa* ParB | This study |
| | pENTR::*Xanthomonas campestris* ParB | This study |
| | pENTR::*Thermus thermophilus* ParB | This study |
| | pENTR::*Bifidobacterium longum* ParB | This study |
| | pENTR::*Mycobacterium tuberculosis* ParB | This study |
| | pENTR::*Streptomyces coelicolor* ParB | This study |
| | pENTR::*Porphyromonas gingivalis* ParB | This study |
| pML477 | Gateway-cloning destination vector for fusion of protein interest to an N-terminally FLAG tag, xylose-inducible promoter, high-copy number plasmid, spectinomycin$^R$ | Gift from Michael Laub |
| pET21b::ParB-(His)6 | overexpression of ParB-(His)$_6$ from an IPTG-inducible T7 promoter | Promega |
| | pET21b::*Caulobacter crescentus*-ParB-(His)$_6$ (WT) | Gift of Christine Jacob Wagner (Lim et al., 2014) |
| | pET21b::*Caulobacter crescentus*-ParB-(His)$_6$ (BDB) | This study |
| | pET21b::*Bacillus subtilis*-Noc-(His)$_6$ | This study |
| | pET21b::*Bacillus subtilis*-Noc-(His)$_6$ (DBD) | This study |
| | pET21b::PtoN1-(His)$_6$ (Q173T179A184G201) | This study |
| | pET21b::PtoN2-(His)$_6$ (R173K179A184G201) | This study |
| | pET21b::PtoN3-(His)$_6$ (R173T179K184G201) | This study |
| | pET21b::PtoN4-(His)$_6$ (R173T179A184R201) | This study |
| | pET21b::PtoN5-(His)$_6$ (Q173K179A184G201) | This study |
| | pET21b::PtoN6-(His)$_6$ (Q173T179K184G201) | This study |
| | pET21b::PtoN7-(His)$_6$ (Q173T179A184R201) | This study |
| | pET21b::PtoN8-(His)$_6$ (R173K179K184G201) | This study |
| | pET21b::PtoN9-(His)$_6$ (R173K179A184R201) | This study |
| | pET21b::PtoN10-(His)$_6$ (R173T179K184R201) | This study |

| | | |
|---|---|---|
| | pET21b::PtoN11-$(His)_6$ (Q173K179K184G201) | This study |
| | pET21b::PtoN12-$(His)_6$ (Q173K179A184R201) | This study |
| | pET21b::PtoN13-$(His)_6$ (Q173T179K184R201) | This study |
| | pET21b::PtoN14-$(His)_6$ (R173K179K184R201) | This study |
| | pET21b::PtoN15-$(His)_6$ (Q173K179K184R201) | This study |
| | pET21b::*Caulobacter crescentus*-ParB-$(His)_6$ (Q162A) | This study |
| | pET21b::*Caulobacter crescentus*-ParB-$(His)_6$ (K171A) | This study |
| | pET21b::*Caulobacter crescentus*-ParB-$(His)_6$ (S172A) | This study |
| | pET21b::*Caulobacter crescentus*-ParB-$(His)_6$ (R173A) | This study |
| | pET21b::*Caulobacter crescentus*-ParB-$(His)_6$ (S174A) | This study |
| | pET21b::*Caulobacter crescentus*-ParB-$(His)_6$ (N178A) | This study |
| | pET21b::*Caulobacter crescentus*-ParB-$(His)_6$ (R181A) | This study |
| | pET21b::*Caulobacter crescentus*-ParB-$(His)_6$ (V226A) | This study |
| | pET21b::*Caulobacter crescentus*-ParB-$(His)_6$ (R227A) | This study |
| | pET21b::*Caulobacter crescentus*-ParB-$(His)_6$ (R234A) | This study |
| | pET21b::*Caulobacter crescentus*-ParB-$(His)_6$ (K245A) | This study |
| | pET21b::*Caulobacter crescentus*-ParB-$(His)_6$ (R248A) | This study |
| | pET21b::ParB-$(His)_6$ chimera 1 | This study |
| | pET21b::ParB-$(His)_6$ chimera 4 | This study |
| | pET21b::*Caulobacter crescentus*-ParB-$(His)_6$ (RTRS) | This study |
| | pET21b::*Caulobacter crescentus*-ParB-$(His)_6$ (RRMT) | This study |
| | pET21b::*Caulobacter crescentus*-ParB-$(His)_6$ (RRLT) | This study |
| | pET21b::*Caulobacter crescentus*-ParB-$(His)_6$ (QRMR) | This study |
| | pET21b::*Caulobacter crescentus*-ParB-$(His)_6$ (QSRR) | This study |
| | pET21b::*Caulobacter crescentus*-ParB-$(His)_6$ (QTNR) | This study |
| | pET21b::*Caulobacter crescentus*-ParB-$(His)_6$ (QRYR) | This study |
| | pET21b::*Caulobacter crescentus*-ParB-$(His)_6$ (QRRR) | This study |
| | pET21b::*Caulobacter crescentus*-ParB-$(His)_6$ (QRKR) | This study |
| | pET21b::*Caulobacter crescentus*-ParB-$(His)_6$ (TEPG) | This study |
| pUT18C::1xFLAG -DEST | Destination vector for Gateway cloning, 1xFLAG tag fused to the N-terminus of protein of interest, carbenicillin$^R$ | This study |
| | pUT18C::1xFLAG-*Bacillus subtilis* Noc | This study |
| | pUT18C::1xFLAG-*Clostridium difficile* Noc | This study |
| | pUT18C::1xFLAG-*Lactobacillus aviarius* Noc | This study |
| | pUT18C::1xFLAG-*Staphylococcus aureus* Noc | This study |
| | pUT18C::1xFLAG-*Bacillus subtilis* ParB | This study |
| | pUT18C::1xFLAG-*Clostridium difficile* ParB | This study |
| | pUT18C::1xFLAG-*Lactobacillus aviarius* ParB | This study |
| | pUT18C::1xFLAG-*Staphylococcus aureus* ParB | This study |
| | pUT18C::1xFLAG-*Caulobacter crescentus* ParB | This study |
| | pUT18C::1xFLAG-*Agrobacterium tumefaciens* ParB | This study |
| | pUT18C::1xFLAG-*Sinorhizobium meliloti* ParB | This study |
| | pUT18C::1xFLAG-*Lawsonia intracellularis* ParB | This study |
| | pUT18C::1xFLAG-*Desulfovibrio vulgaris* ParB | This study |
| | pUT18C::1xFLAG-*Dechloromonas aromatica* ParB | This study |

| | | |
|---|---|---|
| | pUT18C::1xFLAG-*Pseudomonas aeruginosa* ParB | This study |
| | pUT18C::1xFLAG-*Xanthomonas campestris* ParB | This study |
| | pUT18C::1xFLAG-*Thermus thermophilus* ParB | This study |
| | pUT18C::1xFLAG-*Bifidobacterium longum* ParB | This study |
| | pUT18C::1xFLAG-*Mycobacterium tuberculosis* ParB | This study |
| | pUT18C::1xFLAG-*Streptomyces coelicolor* ParB | This study |
| | pUT18C::1xFLAG-*Porphyromonas gingivalis* ParB | This study |
| | pB1H2-w2::*zif268* | (Noyes et al., 2008) via Addgene |
| | pB1H2-w5::*zif268* | (Noyes et al., 2008) via Addgene |
| | pB1H2-w5L::*zif268* | (Noyes et al., 2008) via Addgene |
| | pU3H3::MCS | (Noyes et al., 2008) via Addgene |
| | pU3H3::*zif268* binding site | (Noyes et al., 2008) via Addgene |
| | pB1H2-w2::*Caulobacter* ParB (R104A + Q173K179K184R201) | This study |
| | pB1H2-w5::*Caulobacter* ParB (R104A + Q173K179K184R201) | This study |
| | pB1H2-w5L::*Caulobacter* ParB (R104A + Q173K179K184R201) | This study |
| | pB1H2-w2::*Caulobacter* ParB (R104A + R173A179T184G201) | This study |
| | pU3H3::7bp-*NBS* | This study |
| | pU3H3::14bp-*NBS* | This study |
| | pU3H3::19bp-*NBS* | This study |
| | pU3H3::24bp-*NBS* | This study |
| | pU3H3::19bp-*parS* | This study |

**TABLE S3. PRIMERS**

| Primers | Sequences |
|---|---|
| | **For construction of pUT18C-1xFLAG-DEST** |
| 1934 | acaatttcacacaggaaacagctatggactacaaggacgacgacgacaagggctcg |
| 1935 | acttagttatatcgatgcatcgaaccactttgtacaagaaagctgaacgagaaac |
| 1936 | agctgtttcctgtgtgaaattgttatccgctcacaattc |
| 1937 | tcgatgcatcgatataactaagtaatatggtgcac |
| | **For ChIP-qPCR** |
| ybbD_parSF2 | GTAAGATACCAGGGCAAGG |
| ybbD_parSR2 | TTACTCTGCACAAGCATCA |
| ygcE_NBSF2 | CGCTACGACGCGATGAATAA |
| ygcE_NBSR2 | CTCTGGATCGAATCCACATTCC |
| ompG_F1 | GCGGAGCCTTCAGTCTATTT |
| ompG_R1 | CAAACCACGTTCCACGTTTAC |
| | **For bio-layer interferometry assays** |
| *NBS*_FOR | [Biotin]GGGAtaTTTCCCGGGAAAta |
| *NBS*_REV | taTTTCCCGGGAAAtaTCCC |
| *parS*_FOR | [Biotin]GGGAtgTTTCACGTGAAAca |
| *parS*_REV | tgTTTCACGTGAAAcaTCCC |
| site1_FOR | [Biotin]GGGAtgTTTCTCGAGAAAca |
| site1_REV | tgTTTCTCGAGAAAcaTCCC |
| site10_FOR | [Biotin]GGGAttTTTCgCGcGAAAaa |
| site10_REV | ttTTTCgCGcGAAAaaTCCC |
| site11_FOR | [Biotin]GGGAttTTTCcCGgGAAAaa |
| site11_REV | ttTTTCcCGgGAAAaaTCCC |
| site12_FOR | [Biotin]GGGAtaTTTCACGTGAAAta |
| site12_REV | taTTTCACGTGAAAtaTCCC |
| site13_FOR | [Biotin]GGGAtaTTTCtCGaGAAAta |
| site13_REV | taTTTCtCGaGAAAtaTCCC |
| site14_FOR | [Biotin]GGGAtaTTTCgCGcGAAAta |
| site14_REV | taTTTCgCGcGAAAtaTCCC |
| site2_FOR | [Biotin]GGGAtgTTTCGCGCGAAAca |
| site2_REV | tgTTTCGCGCGAAAcaTCCC |
| site3_FOR | [Biotin]GGGAtgTTTCCCGGGAAAca |
| site3_REV | tgTTTCCCGGGAAAcaTCCC |
| site4_FOR | [Biotin]GGGAtcTTTCACGTGAAAga |
| site4_REV | tcTTTCACGTGAAAgaTCCC |
| site5_FOR | [Biotin]GGGAtcTTTCtCGaGAAAga |
| site5_REV | tcTTTCtCGaGAAAgaTCCC |
| site6_FOR | [Biotin]GGGAtcTTTCgCGcGAAAga |
| site6_REV | tcTTTCgCGcGAAAgaTCCC |
| site7_FOR | [Biotin]GGGAtcTTTCcCGgGAAAga |
| site7_REV | tcTTTCcCGgGAAAgaTCCC |
| site8_FOR | [Biotin]GGGAttTTTCACGTGAAAaa |
| site8_REV | ttTTTCACGTGAAAaaTCCC |
| site9_FOR | [Biotin]GGGAttTTTCtCGaGAAAaa |
| site9_REV | ttTTTCtCGaGAAAaaTCCC |
| | **For the construction of pU3H3-*NBS* and pU3H3-*parS*** |
| *NBS*_anneal_7bp_spacer_F | Ccgggtatttcccggaaataggg |

| | |
|---|---|
| *NBS*_anneal_7bp_spacer_R | aattccctatttcccgggaaatac |
| *NBS*_anneal_14bp_spacer_F | ccgggtatttcccgggaaataggcgcgccg |
| *NBS*_anneal_14bp_spacer_R | aattcggcgcgcctatttcccgggaaatac |
| *NBS*_anneal_19bp_spacer_F | ccgggtatttcccgggaaataggtttcgcgcgccg |
| *NBS*_anneal_19bp_spacer_R | Aattcggcgcgcgaaacctatttcccgggaaatac |
| *NBS*_anneal_24bp_spacer_F | ccgggtatttcccgggaaataggtttcttggcgcgcgccg |
| *NBS*_anneal_24bp_spacer_R | aattcggcgcgcgccaagaa acctatttcccgggaaatac |
| *parS*_anneal_19bp_spacer_F | ccgggtgtttcacgtgaaacaggtttcgcgcgccg |
| *parS*_anneal_19bp_spacer_R | ttaagccgcgcgctttggacaaagtgcactttgtg |
| | **For the construction of AB1157 *ybbD*::*parS ygcE*::*NBS* strain** |
| 1940 | aaatattggagctggattgcctgatgcttgtgcagagtaatgttccacgtggaacaattccggggatccgtcgacctg |
| 1941 | ttgacgacttcgatatgggatagactcttaattcaagcaatgtaggctggagctgcttcg |
| 3139 | ggggaatgtggattcgatccagagctggtcgaatgcgtaatatttcccgggaaataattccggggatccgtcgacctg |
| 3140 | tatgttcaggccgggcagtttcccgcccggccttcctcactgtaggctggagctgcttcgaag |
| | **For generation of Illumina libraries for deep mutational scanning experiments** |
| 4nns_offset_0_F | ACACTCTTTCCCTACACGACGCTCTTCCGATCTgctcaaactattggcaagag |
| 4nns_offset_1_F | ACACTCTTTCCCTACACGACGCTCTTCCGATCTtgctcaaactattggcaagag |
| 4nns_offset_2_F | ACACTCTTTCCCTACACGACGCTCTTCCGATCTttgctcaaactattggcaagag |
| 4nns_offset_3_F | ACACTCTTTCCCTACACGACGCTCTTCCGATCTattgctcaaactattggcaagag |
| 4nns_offset_4_F | ACACTCTTTCCCTACACGACGCTCTTCCGATCTcattgctcaaactattggcaagag |
| 4nns_R | GTGACTGGAGTTCAGACGTGTGCTCTTCCGATCTTTTTGCTAACGCTACGGGATC |
| NEBNext universal primer | AAT GAT ACG GCG ACC ACC GAG ATC TAC ACT CTT TCC CTA CAC GAC GCT CTT CCG ATC-s-T |
| NEBNext Index primer | CAAGCAGAAGACGGCATACGAGATNNNNNNGTGACTGGAGTTCAGACGTGTGCTCTTCCGATC-s-T |
| | **For generation of the deep mutational scanning library** |
| For_B_NNS_HTH | GAGGTACAGTCCTATCTTGTGAGTGGAGAGCTGACAGCGNNSCATGCGCGTGCGATTGCCGCTGC |
| Rev_B_NNS_HTH | GTCCGGCAASNNAAGAAGACGCATSNNATTCGCTACGTGAGASNNACTCTTGCCAATAGTTTGAGC |
| | **For the amplification of the pENTR backbone** |
| pENTR_gibson_backbone_R | ggtgaagggggcggccgcggagcctgc |
| pENTR_gibson_backbone_F | aagggtgggcgcgccgacccagctttcttg |

Nucleotides in red font are spacer bases used to increase the diversity of the Illumina library. Underlined nucleotides are either Illumina library barcodes or NNS bases.

### TABLE S4. X-RAY DATA COLLECTION AND PROCESSING STATISTICS

| **Structure** | **ParB (DBD)-*parS* complex** | **Noc (DBD)-*NBS* complex** |
|---|---|---|
| Data collection | | |
| Diamond Light Source beamline | I04 | I03 |
| Wavelength (Å) | 0.980 | 0.976 |
| Detector | Pilatus 6M-F | Eiger2 XE 16M |
| Resolution range (Å) | 40.12 – 2.40 (2.49 – 2.40) | 72.30 – 2.23 (2.66 – 2.23)[a] |
| Space Group | *C*2 | *C2* |
| Cell parameters (Å/°) | *a* = 122.1, *b* = 40.7, *c* = 94.0, β = 121.4 | *a* = 134.1 *b* = 60.6, *c* = 81.1, β = 116.9 |
| Total no. of measured intensities | 105021 (10942) | 142152 (6183) |
| Unique reflections | 16317 (1662) | 10830 (542) |
| Multiplicity | 6.4 (6.6) | 13.1 (11.4) |
| Mean $I/\sigma(I)$ | 7.0 (2.0) | 9.3 (1.5) |
| Completeness (spherical; %) | 99.7 (99.2) | 38.1 (4.7) |
| Completeness (ellipsoidal; %) | - | 88.4 (57.2) |
| $R_{merge}$[b] | 0.137 (0.801) | 0.108 (0.851) |
| $R_{meas}$[c] | 0.150 (0.869) | 0.112 (0.891) |
| $CC_{½}$[d] | 0.992 (0.850) | 1.000 (0.847) |
| Wilson *B* value (Å$^2$) | 42.1 | 115.7 |
| Refinement | | |
| Resolution range (Å) | 40.12 – 2.40 | 72.30 – 2.23 |
| Reflections: working/free[e] | 15480/826 | 10231/599 |
| $R_{work}$[f] | 0.216 | 0.231 |
| $R_{free}$[f] | 0.232 | 0.279 |
| Ramachandran plot: favoured/allowed/disallowed[g] (%) | 96.5/3.5/0.0 | 95.4/4.6/0.0 |
| R.m.s. bond distance deviation (Å) | 0.003 | 0.002 |
| R.m.s. bond angle deviation (°) | 1.08 | 1.03 |

| | | |
|---|---|---|
| No. of protein residues per chain | 121/140 | 105/116 |
| No. of DNA bases per chain | 20/20 | 22/22 |
| No. of water/glycerol molecules | 82/2 | 0/0 |
| Mean *B* factors: protein/DNA/ water/overall (Å$^2$) | 51/46/38/49 | 155/148/0/154 |
| **PDB accession code** | **6S6H** | **6Y93** |

Values in parentheses are for the outer resolution shell.

[a] After correction by STARANISO to remove poorly measured reflections affected by anisotropy, the ellipsoidal resolutions were:

2.23 Å in direction 0.854 $a^*$ + 0.017 $b^*$ - 0.519 $c^*$

3.83 Å in direction 0.302 $a^*$ + 0.784 $b^*$ + 0.543 $c^*$

4.02 Å in direction 0.130 $a^*$ + 0.911 $b^*$ + 0.391 $c^*$

[b] $R_{merge} = \sum_{hkl} \sum_i |I_i(hkl) - \langle I(hkl)\rangle| / \sum_{hkl} \sum_i I_i(hkl)$.

[c] $R_{meas} = \sum_{hkl} [N/(N-1)]^{1/2} \times \sum_i |I_i(hkl) - \langle I(hkl)\rangle| / \sum_{hkl} \sum_i I_i(hkl)$, where $I_i(hkl)$ is the *i*th observation of reflection *hkl*, $\langle I(hkl)\rangle$ is the weighted average intensity for all observations *i* of reflection *hkl* and *N* is the number of observations of reflection *hkl*.

[d] $CC_{½}$ is the correlation coefficient between symmetry equivalent intensities from random halves of the dataset.

[e] The dataset was split into "working" and "free" sets consisting of 95 and 5% of the data respectively. The free set was not used for refinement.

[f] The R-factors $R_{work}$ and $R_{free}$ are calculated as follows: $R = \sum(| F_{obs} - F_{calc} |)/\sum| F_{obs} |$, where $F_{obs}$ and $F_{calc}$ are the observed and calculated structure factor amplitudes, respectively.

[g] As calculated using MolProbity (Chen et al., 2010).

### TABLE S5. DEEP MUTATIONAL SCAN AND ChIP-Seq

| **Deep mutational scanning libraries** | **GEO** |
|---|---|
| Pre-selection library, replicate 1 | This study ( GSE129285) |
| Pre-selection library, replicate 2 | This study ( GSE129285) |
| Pre-selection library, replicate 3 | This study ( GSE129285) |
| Post-selection library, selection for *parS*-binding capability, replicate 1 | This study ( GSE129285) |
| Post-selection library, selection for *parS* -binding capability, replicate 2 | This study ( GSE129285) |
| Post-selection library, selection for *parS* -binding capability, replicate 3 | This study ( GSE129285) |
| Post-selection library, selection for *NBS*-binding capability, replicate 1 | This study ( GSE129285) |
| Post-selection library, selection for *NBS* -binding capability, replicate 2 | This study ( GSE129285) |
| Post-selection library, selection for *NBS* -binding capability, replicate 3 | This study ( GSE129285) |
| **ChIP-seq datasets** | GEO |
| TLE3000+pUT18C::1xFLAG-Bacillus subtilis Noc, fixation with 1% formaldehyde, α-FLAG antibody (Sigma), ChIP fraction | This study ( GSE129285) |
| TLE3000+pUT18C::1xFLAG-Clostridium difficile Noc, fixation with 1% formaldehyde, α-FLAG antibody (Sigma), ChIP fraction | This study ( GSE129285) |
| TLE3000+pUT18C::1xFLAG-Lactobacillus aviarius Noc, fixation with 1% formaldehyde, α-FLAG antibody (Sigma), ChIP fraction | This study ( GSE129285) |
| TLE3000+pUT18C::1xFLAG-Staphylococcus aureus Noc, fixation with 1% formaldehyde, α-FLAG antibody (Sigma), ChIP fraction | This study ( GSE129285) |
| TLE3000+pUT18C::1xFLAG-Bacillus subtilis ParB, fixation with 1% formaldehyde, α-FLAG antibody (Sigma), ChIP fraction | This study ( GSE129285) |
| TLE3000+pUT18C::1xFLAG-Clostridium difficile ParB, fixation with 1% formaldehyde, α-FLAG antibody (Sigma), ChIP fraction | This study ( GSE129285) |
| TLE3000+pUT18C::1xFLAG-Lactobacillus aviarius ParB, fixation with 1% formaldehyde, α-FLAG antibody (Sigma), ChIP fraction | This study ( GSE129285) |
| TLE3000+pUT18C::1xFLAG-Staphylococcus aureus ParB, fixation with 1% formaldehyde, α-FLAG antibody (Sigma), ChIP fraction | This study ( GSE129285) |
| TLE3000+pUT18C::1xFLAG-Caulobacter crescentus ParB, fixation with 1% formaldehyde, α-FLAG antibody (Sigma), ChIP fraction | This study ( GSE129285) |
| TLE3000+pUT18C::1xFLAG-Agrobacterium tumefaciens ParB, fixation with 1% formaldehyde, α-FLAG antibody (Sigma), ChIP fraction | This study ( GSE129285) |
| TLE3000+pUT18C::1xFLAG-Sinorhizobium meliloti ParB, fixation with 1% formaldehyde, α-FLAG antibody (Sigma), ChIP fraction | This study ( GSE129285) |
| TLE3000+pUT18C::1xFLAG-Lawsonia intracellularis ParB, fixation with 1% formaldehyde, α-FLAG antibody (Sigma), ChIP fraction | This study ( GSE129285) |
| TLE3000+pUT18C::1xFLAG-Desulfovibrio vulgaris ParB, fixation with 1% formaldehyde, α-FLAG antibody (Sigma), ChIP fraction | This study ( GSE129285) |
| TLE3000+pUT18C::1xFLAG-Dechloromonas aromatica ParB, fixation with 1% formaldehyde, α-FLAG antibody (Sigma), ChIP fraction | This study ( GSE129285) |
| TLE3000+pUT18C::1xFLAG-Pseudomonas aeruginosa ParB, fixation with 1% formaldehyde, α-FLAG antibody (Sigma), ChIP fraction | This study ( GSE129285) |
| TLE3000+pUT18C::1xFLAG-Xanthomonas campestris ParB, fixation with 1% formaldehyde, α-FLAG antibody (Sigma), ChIP fraction | This study ( GSE129285) |
| TLE3000+pUT18C::1xFLAG-Thermus thermophilus ParB, fixation with 1% formaldehyde, α-FLAG antibody (Sigma), ChIP fraction | This study ( GSE129285) |
| TLE3000+pUT18C::1xFLAG-Bifidobacterium longum ParB, fixation with 1% formaldehyde, α-FLAG antibody (Sigma), ChIP fraction | This study ( GSE129285) |

| | |
|---|---|
| TLE3000+pUT18C::1xFLAG-Mycobacterium tuberculosis ParB, fixation with 1% formaldehyde, α-FLAG antibody (Sigma), ChIP fraction | This study ( GSE129285) |
| TLE3000+pUT18C::1xFLAG-Streptomyces coelicolor ParB, fixation with 1% formaldehyde, α-FLAG antibody (Sigma), ChIP fraction | This study ( GSE129285) |
| TLE3000+pUT18C::1xFLAG-Porphyromonas gingivalis ParB, fixation with 1% formaldehyde, α-FLAG antibody (Sigma), ChIP fraction | This study ( GSE129285) |
